# Extracellular glutamate and GABA transients at the transition from interictal spiking to seizures

**DOI:** 10.1101/2020.11.13.381707

**Authors:** Yoshiteru Shimoda, Marco Leite, Robert T Graham, Jonathan S Marvin, Jeremy Hasseman, Ilya Kolb, Loren L Looger, Vincent Magloire, Dimitri M Kullmann

**Author notes:** These authors contributed equally (Vincent Magloire, Dimitri M Kullmann). Department of Neurosurgery, Tohoku University Graduate School of Medicine, Sendai 980-8575, Japan (Yoshiteru Shimoda). University of California San Diego, 9500 Gilman Drive #, La Jolla CA 92093, USA. Author contributions: Y.S., V.M. and D.M.K. conceived the experiments. Y.S and V.M. performed the experiments. Y.S., V.M., R.T.G. and M.L. analysed the data. M.L. performed the simulations. J.S.M., J.H., I.K. and L.L.L. provided the iGABASnFR constructs. V.M., Y.S. and D.M.K. wrote the manuscript, which was revised by all the authors. D.M.K. and V.M. secured funding.

## Abstract

Focal epilepsy is associated with intermittent brief population discharges (interictal spikes), which resemble sentinel spikes that often occur at the onset of seizures. Why interictal spikes self-terminate whilst seizures persist and propagate is incompletely understood. We used fluorescent glutamate and GABA sensors in an awake rodent model of neocortical seizures to resolve the spatiotemporal evolution of both neurotransmitters in the extracellular space. Interictal spikes were accompanied by brief glutamate transients which were maximal at the initiation site and rapidly propagated centrifugally. GABA transients lasted longer than glutamate transients and were maximal ∼1.5 mm from the focus where they propagated centripetally. Prior to seizure initiation GABA transients were attenuated, whilst glutamate transients increased, consistent with a progressive failure of local inhibitory restraint. As seizures increased in frequency, there was a gradual increase in the spatial extent of spike-associated glutamate transients associated with interictal spikes. Neurotransmitter imaging thus reveals a progressive collapse of an annulus of feed-forward GABA release, allowing seizures to escape from local inhibitory restraint.

## Introduction

Epilepsy is characterized by a propensity for seizures, which typically occur only intermittently but can have devastating consequences for the patient if they cannot be suppressed by pharmacological or surgical intervention. Many types of seizures are recognized, with various clinical manifestations, electro-encephalographic (EEG) correlates and aetiologies. Nevertheless, a very common pattern is for seizures to arise from a specific cortical region, and to spread to involve local networks or to generalize to both hemispheres. Such focal or partial-onset epilepsy is very often accompanied by interictal discharges (IIDs), which can be used clinically to help localize the seizure onset zone ^1^. IIDs (also known as interictal spikes) are electrographically similar to large-amplitude discharges that often occur at the start of a seizure (so-called sentinel spikes ^2^). The mechanisms underlying interictal spike generation and propagation are incompletely understood, as are the reasons why spikes can sometimes proceed to seizures. Interictal spikes are dominated by GABAergic signalling, consistent with powerful feed-forward inhibition confining the spatial and temporal extent of hypersynchronous firing of principal neurons ^3–6^. Collapse of this inhibitory restraint provides a parsimonious explanation why seizures occur intermittently. This principle is supported by *in vitro* models, as well as *in vivo* electrical recordings in both experimental models and patients with refractory epilepsy ^5–10^. It is also consistent with the spatiotemporal evolution of neuronal activity measured using fluorescent calcium indicators in awake head-fixed rodents, where interictal spikes and seizures are initially indistinguishable ^11^.

How feed-forward inhibition collapses remains unclear. Several possible mechanisms have been proposed. These include chloride accumulation in principal neurons upon intense activation of GABA_A_ receptors, which could in turn attenuate the inhibitory effect of these receptors and even give rise to GABA-mediated depolarization ^12, 13^, facilitating the recruitment of principal neurons. Chloride build-up could also lead to extracellular potassium accumulation through activation of the Cl^-^/K^+^ symporter KCC2 ^14–16^. Alternatively, collapse of an inhibitory restraint could occur as a result of a failure of GABA release, for instance because of vesicle depletion, presynaptic inhibition of GABAergic terminals ^17, 18^, or because intense depolarization of interneurons prevents them from firing (depolarization block) ^19, 20^. Evidence has been put forward in support of all of the above hypotheses, and they are not mutually exclusive, especially when the diversity of interneurons is taken into account ^21^. However, much of the available evidence comes from highly reduced preparations of debatable relevance to seizure mechanisms *in vivo*. Furthermore, experimental manipulations that interfere with different aspects of inhibitory signalling to investigate how inhibition collapses are themselves pro-epileptic.

Imaging epileptiform activity in awake head-fixed animals offers several advantages, including preservation of an intact circuitry and neurochemical milieu, and the ability to monitor activity at a scale allowing the early transition of local interictal spikes to propagating seizures to be detected. Hitherto, imaging of seizures *in vivo* has mainly relied on calcium-sensitive fluorescent reporters of neuronal activity ^11, 22–24^, which are sensitive but have not been exploited extensively in the context of seizure activity to distinguish between principal neurons and interneuron subtypes. Here we take an alternative approach to monitoring the transition from interictal to seizure activity, by imaging extracellular glutamate and GABA transients in the vicinity of a focus elicited by intracortical chemoconvulsant injection ^25^. Indeed, this approach has the potential to uncover aspects of the initiation and propagation of seizures that cannot be detected electrographically. We show striking differences in the spatiotemporal evolution of the two neurotransmitters, and provide evidence that seizures are heralded by a shift in the relative balance of glutamate and GABA transients. The data argue that the collapse of inhibitory restraint is at least partly due to a failure of GABA release.

## Results

### Extracellular GABA and glutamate imaging in acute chemoconvulsant epilepsy models

We imaged extracellular glutamate and GABA using variants of fluorescent reporters based on bacterial periplasmic binding proteins ^25, 26^ expressed in the visual cortex of adult mice by injection of adeno-associated viral vectors (AAVs). Both iGluSnFR and iGABASnFR incorporate the PDGFR transmembrane domain, and their fluorescence responds to both synaptic and extrasynaptic neurotransmitter kinetics. Micron-scale imaging was not attempted, and instead we used spiral line scans across most of the field of view to characterize the time course of the average extracellular concentrations of GABA and glutamate approximately 100 µm below the pial surface (**Fig. 1A–E**). On the day of the experiment, the GABA_A_ receptor antagonist picrotoxin (PTX), or the cholinergic agonist pilocarpine (Pilo), was microinjected into cortical layer 5 (at least 1mm caudal to the imaging window, **Fig. 1A, C**). By recording the electrocorticogram (ECoG) *via* an electrode implanted in the vicinity of the injection site, we obtained concurrent electrophysiological data on the evolution of epileptiform activity, which typically transitioned over 10 – 20 minutes from IIDs to brief seizures lasting a few seconds (median ± median absolute deviation: 5.64 ± 0.25 s), each of which was heralded by a sentinel spike (**Fig. 1F**).

**Figure 1:**
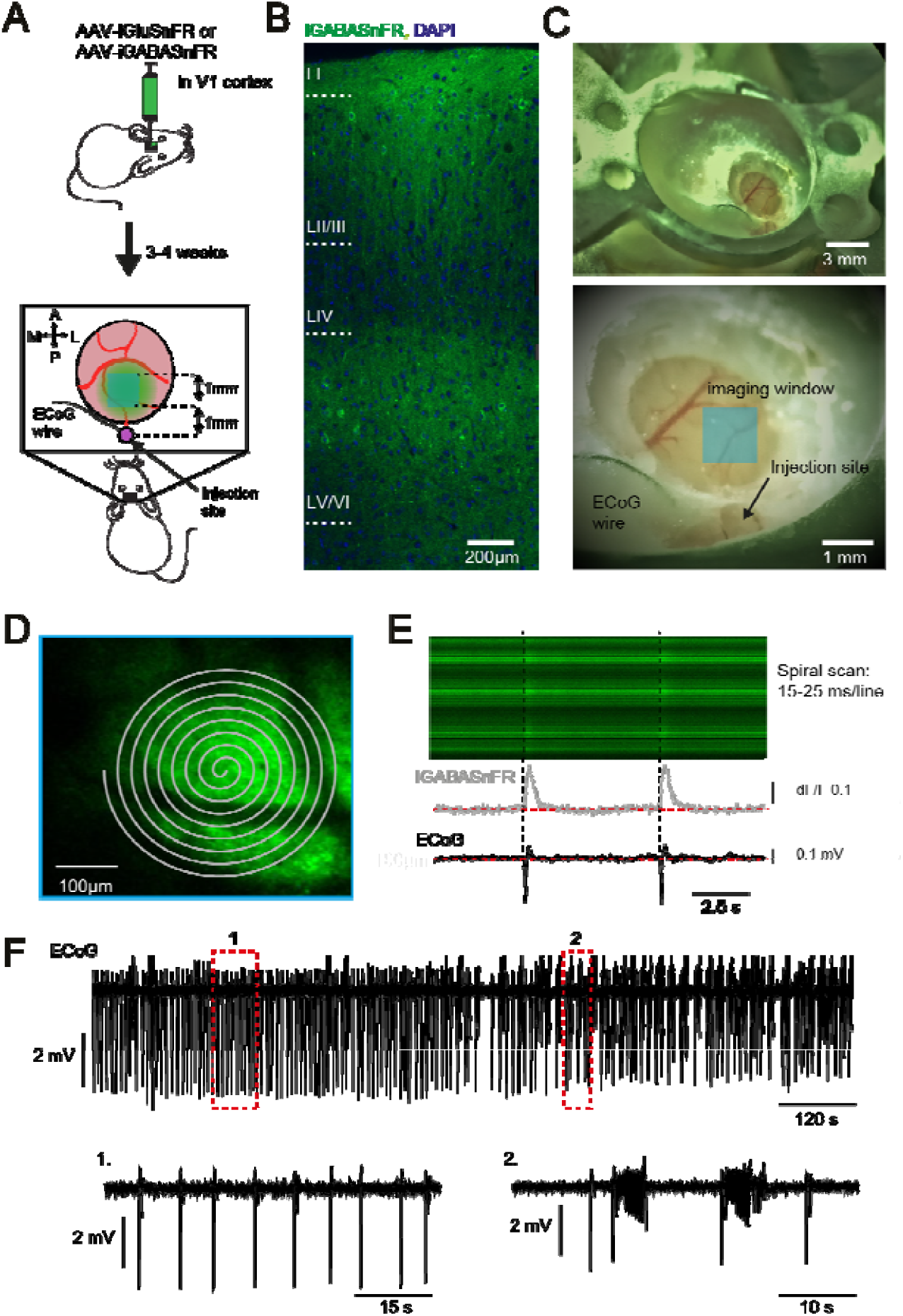
Protocol to characterize spatiotemporal neurotransmitter profiles during epileptic activity. (**A**) Experimental design. (**B**) Expression of iGABASnFR (green) and DAPI staining (blue) across layers of the visual cortex. The imaging plane was ∼100µm below the pia. (**C**) Photograph of the headpiece and craniotomy. The site of chemoconvulsant injection was ∼1mm from the imaging window (blue square) and the ECoG electrode was positioned nearby. (**D**) Frame scan showing iGABASnFR fluorescence, with the spiral line scan trajectory superimposed. (**E**) Aligned linescan (top), average iGABASnFR signal and ECoG trace during interictal spiking. (**F**) Example of ECoG recording of epileptiform activity, initially dominated by IIDs (red rectangle 1 expanded below), which gave way to intermittent seizures lasting several seconds and heralded by sentinel spikes (red rectangle 2).

### Temporal profiles of glutamate and GABA during IIDs

During the initial interictal spiking period, IIDs were accompanied by large amplitude iGluSnFR and iGABASnFR fluorescence transients (**Fig. 2A, B**). The rise-time of iGABASnFR fluorescence transients (measured from 10% to peak) following PTX application was approximately 2-fold slower than that of iGluSnFR transients (iGluSnFR: 63 ± 6 ms, n=14 mice, average ± s.e.m.; iGABASnFR: 134 ± 10 ms, n= 11 mice, p=2.15×10^-6^, unpaired t-test). A qualitatively similar difference in rise-times was seen in response to Pilo application although it fell short of significance (iGluSnFR: 99 ± 18 ms, n=5 mice, average ± SEM; iGABASnFR: 139 ± 9 ms, n= 4 mice, p=0.11, unpaired t-test). The difference in rise-times between iGABASnFR and iGluSnFR signals is consistent with the slower binding and activation kinetics of iGABASnFR than iGluSnFR ^25^. GABA sensor fluorescence transients also decayed more slowly than glutamate sensor transients in response to both chemoconvulsants (monoexponential *τ* fitted from peak to 1 s after the peak: PTX, iGluSnFR: 95 ± 9 ms, n= 14 mice; iGABASnFR: 264 ± 27 ms, n = 11 mice; p=1.25×10^-6^ unpaired t-test, **Fig. 2C;** Pilo, iGluSnFR: 85 ± 7 ms, n= 5 mice; iGABASnFR: 175 ± 27 ms, n = 4 mice; p=0.009 unpaired t-test, **Fig. 2E**). As an alternative measure of kinetics, we integrated the decaying phase of the fluorescence waveforms (area under the curve, AUC) and divided by the peak amplitude. This measure confirmed that the fluorescence decay kinetics were slower for the GABA than the glutamate sensor (PTX: iGluSnFR: 704 ± 96 ms; iGABASnFR: 1450 ± 37 ms; p=1.01×10^-6^ unpaired t-test, **Fig. 2D;** Pilo: iGluSnFR: 514 ± 79 ms; iGABASnFR: 722 ± 30 ms; p=0.06 unpaired t-test, **Fig. 2F**). This difference in decay kinetics is unlikely to be due to an intrinsic difference in sensor off-rates because the decay time-constants previously estimated *in vitro* in response to electrical stimulation of neuronal cultures were similar for both sensors ^25, 26^, and similar to the decay for iGluSnFR during interictal spiking (*τ* < 100 ms). The slower decay of iGABASnFR transients *in vivo* implies that GABA released during IIDs is cleared from the extracellular space more slowly than glutamate (It does not necessarily indicate that more GABA than glutamate is released because GABA transporters are expressed at a lower density than glutamate transporters ^27–29^).

**Figure 2:**
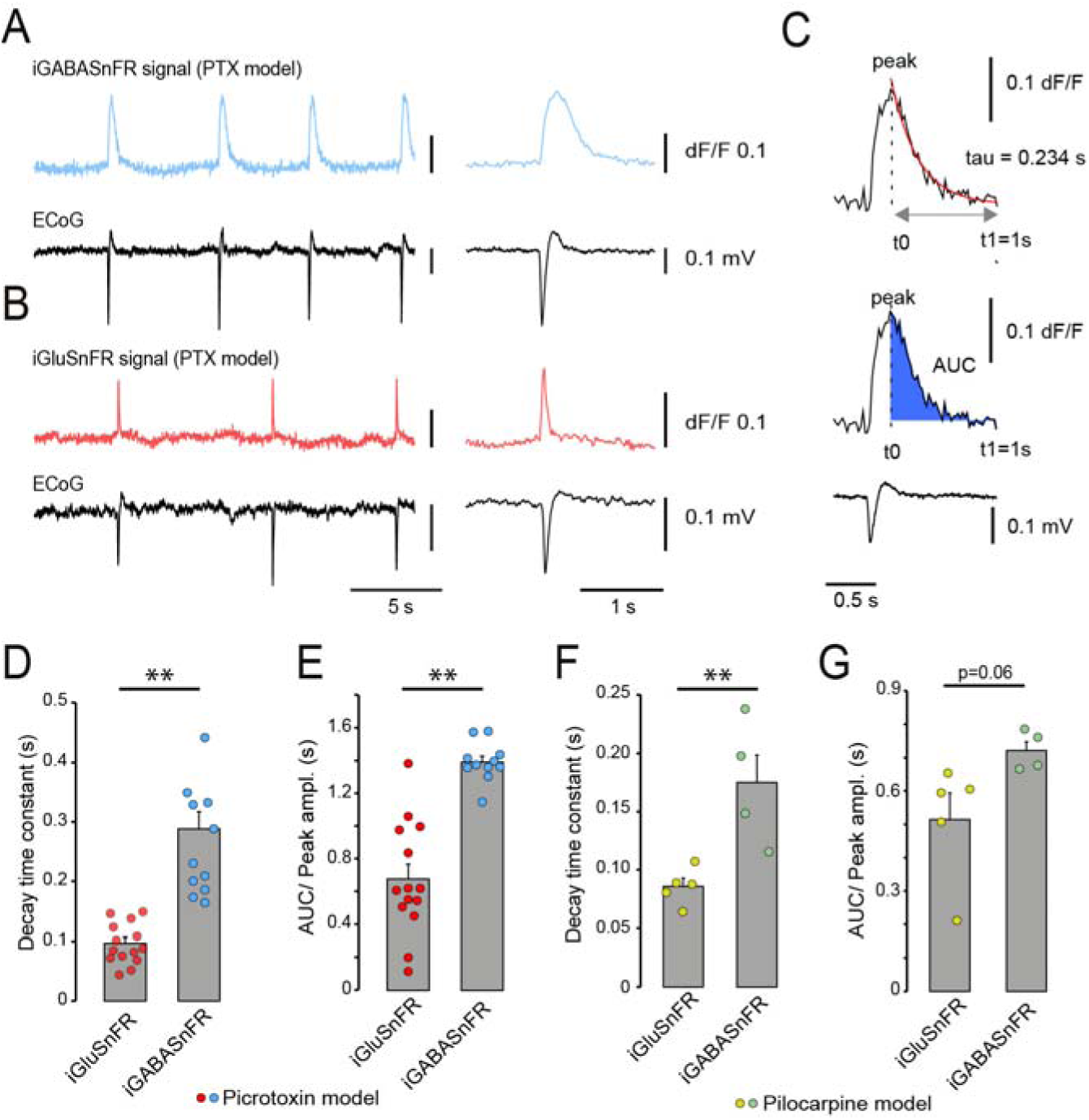
Extracellular GABA persists for longer than glutamate during IIDs in picrotoxin and pilocarpine acute epilepsy models. (**A**) iGluSnFR fluorescence and simultaneous ECoG. (**B**) Same for iGABASnFR. (**C**) Measurements of decay time constant and the area under the curve from peak to 1s later (example trace from one iGABASnFR experiment). (**D**) Average decay time constant of the glutamate (red symbols) and GABA (blue symbols) transients following PTX application in 14 mice expressing iGluSnFR and 11 expressing iGABASnFR. (**E**) Average area under the curve divided by peak amplitude (AUC/Peak ampl.) in the same experiments. (**F**) Average decay time constant of the glutamate (yellow symbols) and GABA (green symbols) transients following Pilo application in 5 mice expressing iGluSnFR and 4 expressing iGABASnFR. (**G**) AUC/Peak ampl. in the same experiments. Error bars in (D, E) and (F, G) correspond to s.e.m.. **p<0.01.

The data summarized above were obtained in experiments where either iGluSnFR or iGABASnFR was expressed and imaged separately. The availability of spectrally shifted variants of iGluSnFR ^30^ permits both neurotransmitters to be imaged simultaneously. We estimated the unmixing coefficients required to isolate the fluorescence of SF-Venus-iGluSnFR.A184V (yellow) and GFP-iGABASnFR (green) whilst imaging with three different emission filters (see Materials and Methods and **Fig. S1**). Simultaneous imaging of both sensors confirmed a slower decay of GABA than glutamate fluorescence in all 3 animals imaged (PTX model, **Fig. S1**).

Taken together, these observations imply that IIDs are accompanied by substantial release of both glutamate and GABA, and that GABA persists in the extracellular space for longer than glutamate. When comparing between the chemoconvulsants, the iGABASnFR signal persisted for longer following PTX than Pilo application (**Fig. S2**). There was also a non-significant trend for iGluSnFR signals to decay more slowly following Pilo than PTX. A possible explanation is that Pilo diffused further from the application site. Apart from this difference, the PTX and Pilo models yielded very similar data. The consistency of results obtained with two chemoconvulsant models argues that imaging GABA and glutamate uncovers fundamental mechanisms relating to IIDs.

### Spatial profiles of glutamate and GABA during IIDs

Previous Ca^2+^ imaging experiments using either pilocarpine or picrotoxin microinjection to evoke epileptiform activity showed that interictal events self-terminate and remain confined to a focus of radius ∼1 mm, whilst other, initially indistinguishable, electrographic events turn into seizures that invade adjacent cortical territories ^11^. We asked how neurotransmitter transients during IIDs relate to the distance from the ictal focus in the critical transition zone spanning 1 – 2mm from the site of chemoconvulsant application. Spiral scans were used to image iGluSnFR or iGABASnFR repeatedly at different locations within this region (**Fig. S3**). In all but one experiment, the peak intensity of the iGluSnFR fluorescence decreased with distance from the injection site. In striking contrast, the peak of iGABASnFR fluorescence intensity was maximal at intermediate distances from the focus (**Fig. 3A–D**). This pattern held whether pilocarpine or picrotoxin was used to elicit IIDs (**Fig. 3C,D**). Expressed as a distance-weighted average within the imaging window, the GABA fluorescence transient was significantly further from the focus than the glutamate signal (pooled data from PTX and Pilo models: iGluSnFR: 1.36 ± 0.02 mm, n= 12 mice; iGABASnFR: 1.48 ± 0.03 mm, n= 8 mice, p=0.005, unpaired t-test, **Fig. 3E**; because imaging was confined to the interval 1 – 2mm, the difference between the weighted averages underestimates the spatial separation between the peaks of glutamate and GABA). These findings are consistent with GABA release acting as an inhibitory halo around a hyperexcitable focus, analogous with feed-forward inhibition ahead of ictal wavefronts in epileptic patients and animal models ^7, 9^.

**Figure 3:**
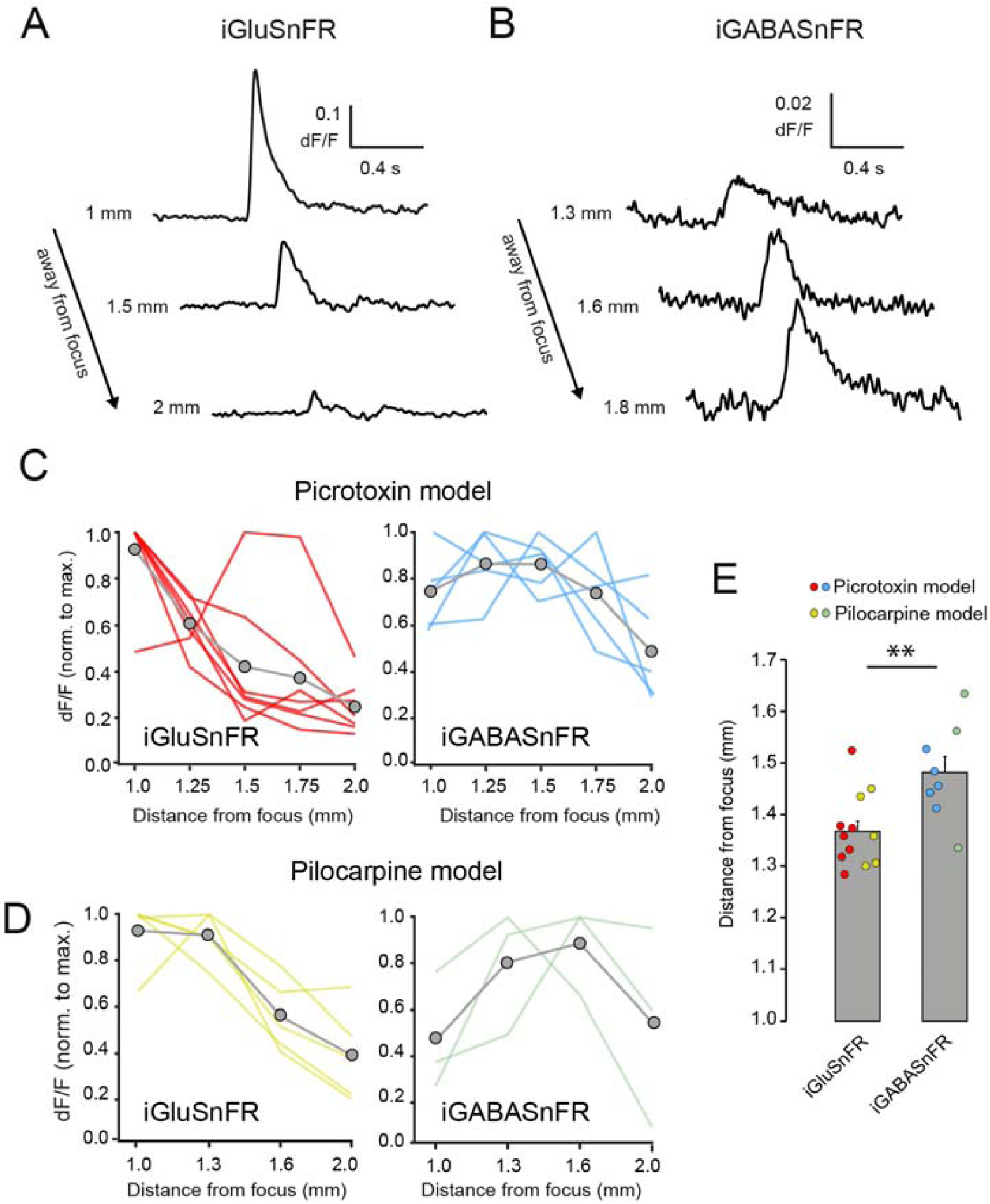
Contrasting spatial profile of glutamate and GABA during IIDs. (**A**) Example iGluSnFR fluorescence transients at different distances from the focus during interictal spiking in one experiment (PTX model). (**B**) Example iGABASnFR transients (PTX model). (**C**) The iGluSnFR peak transient amplitude (red lines) diminished with distance from the focus, whilst the iGABASnFR transient (blue lines) was maximal ∼1.5 mm away (light coloured lines: individual experiments, grey line: average). In each experiment the spatial profile was normalized to the maximal fluorescence transient. (**D**) similar spatial profiles of glutamate and GABA transients were observed with pilocarpine injection (yellow lines: iGluSnFR, green lines: iGABASnFR). (**E**) Peak-weighted average distance (∑((dF/F*distance)/∑(dF/F)) of iGluSnFR and iGABASnFR transients within the imaging interval between 1 mm and 2 mm (n=7 and 5 mice for iGluSnFR using PTX (red symbols) and Pilo (yellow symbols), and 5 and 3 mice for iGABASnFR respectively (blue symbols: PTX, green symbols: Pilo)). Error bars in (E) correspond to s.e.m.. **p<0.01.

We estimated the direction of propagation of the waves of glutamate and GABA during IIDs by calculating the time taken to reach 50% of the maximal fluorescence intensity at different regions in the imaging plane, and then relating their locations to a vector that was rotated through 360*°*. The vector angle that yielded the steepest correlation between time-lag and distance along the vector defined the direction of propagation of the neurotransmitter wave (**Fig. S4**). This procedure also yielded an estimate of the speed of propagation of the interictal fluorescence transients. For iGluSnFR fluorescence transients, the wave propagated away from the focus, with a velocity of 55.9 ± 9.9 mm/s (no difference was found between PTX and Pilo; pooled data n=13 mice). For iGABASnFR, in contrast, the wave propagated towards the focus, at a significantly slower velocity (pooled data: 23.5 ± 7.2 mm/s, n=9 mice; p=0.024, unpaired t-test, **Fig. 4**). This striking difference further supports the view that GABA release acts to constrain the spatial extent of IIDs.

**Figure 4:**
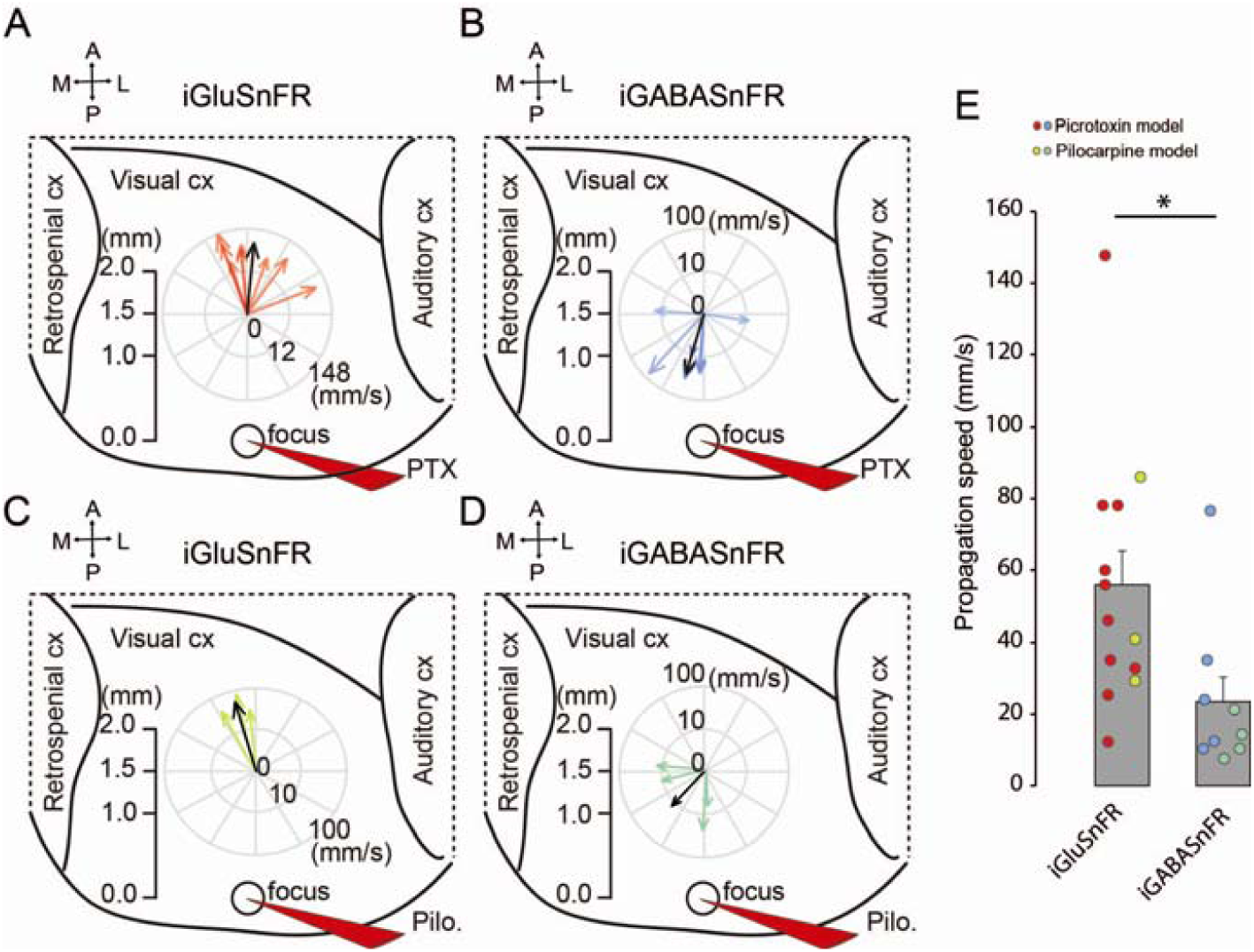
Glutamate and GABA waves propagate in opposite directions. (**A**) Propagation direction and velocity of the iGluSnFR signal in 10 animals (red arrows, average vector shown as black arrow) during IIDs elicited by PTX, imaged ∼1.5 mm away from the focus. (**B**) Same but for iGABASnFR fluorescence transients (blue arrows). (**C, D**) Experiments using Pilo instead of PTX, showing glutamate (yellow arrows) and GABA fluorescent signal propagation (green arrows). Glutamate and GABA waves travelled in opposite directions and at different velocities (note the logarithmic radial velocity scales in (A, B) and (C, D)). (**E**) Average propagation speed was significantly slower for the GABA waves (n=10 (red symbols: PTX) and 3 (yellow symbols: Pilo) mice for iGluSnFR and n=5 (blue symbols: PTX) and 4 (green symbols: Pilo) mice for iGABASnFR). Error bars in (E) correspond to s.e.m.. *p<0.05.

### Interictal discharges gradually give way to seizures

The chemoconvulsant models studied here are characterized by a gradual increase in the frequency and duration of seizures, and a decrease in the occurrence of IIDs (PTX model **Fig. 1F** and see ref. ^31^ for Pilo). Previous *in vitro* studies have prompted the view that a progressive loss of preictal inhibitory barrages occurs over hundreds of seconds, facilitating the propagation of electrographic seizures ^6, 10^. We asked two questions. First, do iGABASnFR and iGluSnFR fluorescence transients shed light on why some events terminate immediately (IIDs) whilst others turn into seizures? And second, how do the spatial profiles of GABA and glutamate transients accompanying epileptiform events evolve on a timescale of minutes? To address these questions, we focused on the PTX model both because it was more reproducible than the Pilo model, with a highly stereotypical transition from interictal spikes to seizures, and, as discussed above, because the shorter IID-associated iGABASnFR signals in the Pilo model raise the possibility that pilocarpine was not completely confined to the application site.

### Partial collapse of GABA release, accompanied by an increase of glutamate release, at seizure onset

What distinguishes IIDs from sentinel spikes that evolve into an ictus? We focused on iGABASnFR and iGluSnFR fluorescence transients imaged ∼1.5 – 2.5 mm from the focus, where maximal GABA responses were detected during IIDs (**Fig. 3C, D**). This coincides with the shortest distance at which cortical territory invasion can be detected with Ca^2+^ imaging, distinguishing seizures from IIDs ^11^. GABA fluorescence transients thus measured were smaller when accompanying sentinel spikes than when associated with IIDs (sentinel spike to IID ratio: 0.87 ± 0.04, p=0.013, paired t-test, n=11 mice; **Fig. 5A, B**). In contrast, iGluSnFR transients increased over two-fold at seizure onset relative to IIDs (ratio: 2.59 ± 0.46, p=0.019, paired t-test, n=6 mice; **Fig. 5C, D**).

**Figure 5:**
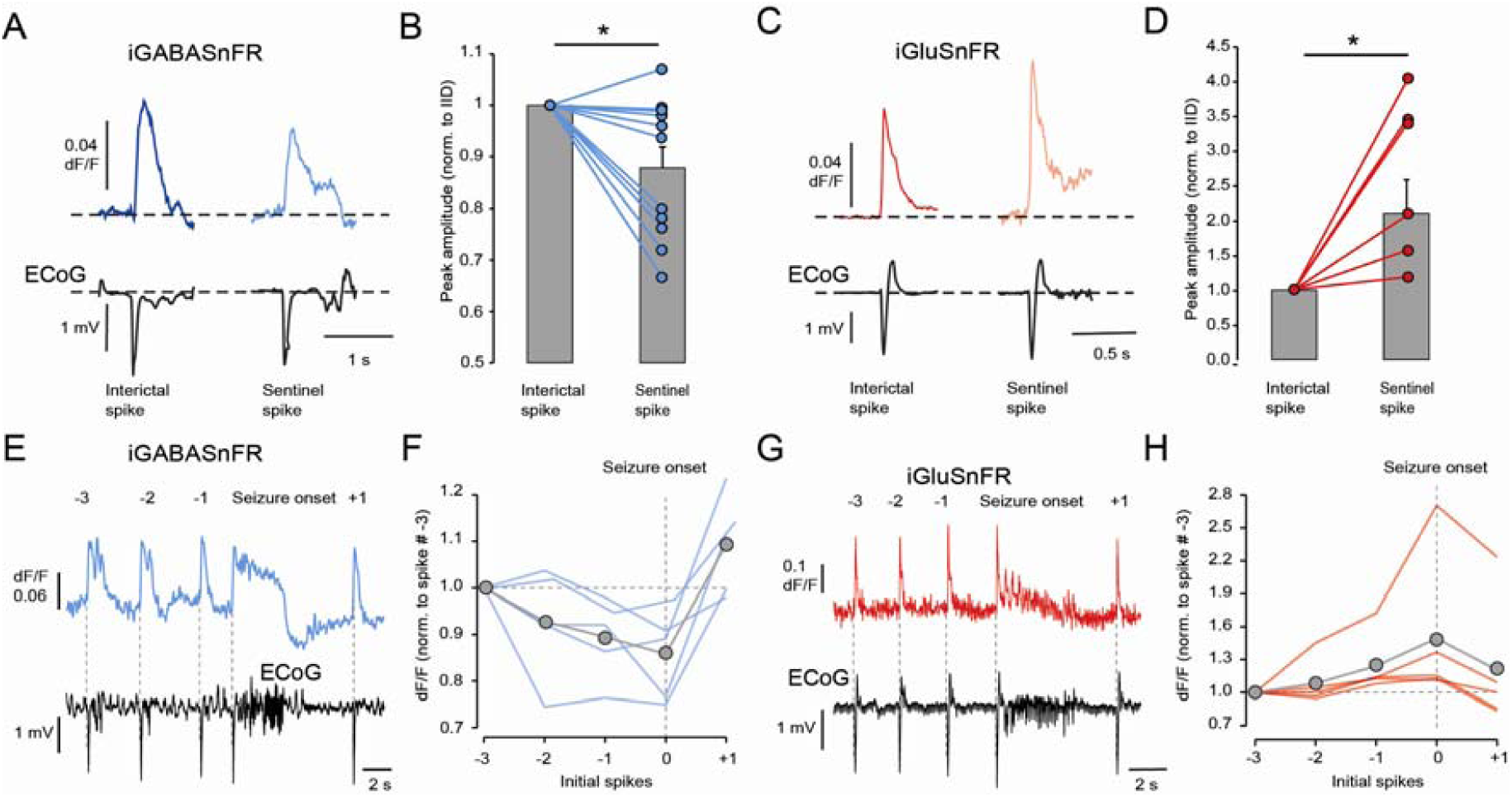
Rapid changes in iGABASnFR and iGluSnFR fluorescence at the transition from interictal spiking to seizures in PTX model. (**A**) Average iGABASnFR fluorescence transient and ECoG trace during interictal spikes (dark blue, n=58 events) and sentinel spikes at seizure onset (light blue, n=25 events; data from one representative experiment). (**B**) Average peak amplitude of iGABASnFR signal accompanying the sentinel spike heralding a seizure, normalized by the average IID-associated signal (n=11 mice). (**C**) same as (A) but for iGluSnFR signal (dark red, n=126 IIDs; light red, n=8 seizures). (**D**) Average peak amplitude of iGluSnFR signal accompanying sentinel spikes normalized by IIDs (n=6 mice). (**E**) iGABASnFR fluorescence trace and ECoG showing 3 IIDs before seizure onset, and one after (numbered). (**F**) Normalized iGABASnFR fluorescence amplitude corresponding to each interictal spike and the sentinel spike, normalized to spike # −3. (Thin coloured lines represent the mean of 3 sequences for each animal, and the thick black line shows the average from all 5 mice). (**G, H**) same as (**E**) and (**F**) respectively but for the glutamate transients. Error bars in (B) and (D) correspond to s.e.m.. *p<0.05.

In some animals, we were able to follow the evolution of GABA and glutamate transients for several consecutive IIDs before and after a seizure. We restricted attention to sequences consisting of 3 interictal spikes before, and one after, a sentinel spike and seizure. This revealed a consistent decrease in iGABASnFR transient amplitude *before* seizure onset (dF/F peak: F[1,4]=1131.94, p=4.65×10^-6^, n=5 mice, one-way RM-ANOVA), which was accompanied by a progressive increase in iGluSnFR signal (dF/F peak: F[1,4]=59.59, p=0.001, n=5 mice, one-way RM-ANOVA), and a recovery after the seizure (3 sequences per mice, 5 mice for each condition, **Fig. 5E-H**).

Taken together, these findings imply that seizure onset is heralded by an incipient failure of GABA release with successive IIDs, and a gradual increase in the ratio of glutamate to GABA release, providing a parsimonious explanation for an escape of epileptiform activity from surround inhibition.

### Slow erosion of GABA release accompanies progressive invasion of the neighbouring cortical territory

To investigate the slow evolution of GABA and glutamate transients over several tens of minutes, we divided the period during which seizures occur into early (<500s from first electrographic seizure) and late (>500s from first seizure) epochs. During this progression the fraction of time spent in a seizure gradually increased (**Fig. S5**). The whole experiment was thus divided into initial interictal spiking (IIS), and early and late seizure activity (ESA and LSA, respectively) (**Fig. 6A–C**). At any stage of the gradual evolution through ESA and LSA (**Fig. 6A**), the spatial profiles of GABA or glutamate transients accompanying sentinel spikes were similar to those corresponding to interspersed IIDs. We therefore averaged sentinel spikes and IIDs together (‘spikes’) in order to study the gradual evolution of the amplitude and shape of glutamate and GABA profiles (**Fig. S5**). Spike-associated iGluSnFR transients increased significantly as the cortical activity gradually evolved from IIS through ESA to LSA (Peak amplitude: F[1,5]=64.48, p=4.84×10^-4^, n=6 mice, RM-ANOVA, **Fig. 6D**). The increase in glutamate transients was most marked as IIS gave way to ESA (Peak amplitude ESA/IIS: 2.06 ± 0.33, p=0.021; LSA/IIS: 2.66 ± 0.39, p=0.001, n=6 mice, Bonferroni test, **Fig. 6A**). In addition, the inverse relationship between amplitude and distance seen for iGluSnFR fluorescence during IIS (**Fig. 3**) disappeared (**Fig. 6E**). The greater invasion of the cortex implies a collapse of the spatial constraint of glutamatergic signalling. Interestingly, this pattern was observed whether restricting attention to the sentinel spikes of individual brief seizures or to the interictal spikes that continued to occur in between seizures (**Fig. S5**). In contrast to glutamate transients, the iGABASnFR signal diminished as IIS gave way to ESA and LSA (Peak amplitude: F[1,6]=239.98, p=4.57×10^-6^, n=7 mice, RM-ANOVA; Peak amplitude: ESA/IIS: 0.82 ± 0.08, p=0.19; LSA/IIS: 0.75 ± 0.10, p=0.04, n=7 mice, Bonferroni test, **Fig. 6F**). The relationship between peak amplitude and distance also showed a tendency to flatten out (**Fig. 6G**).

**Figure 6:**
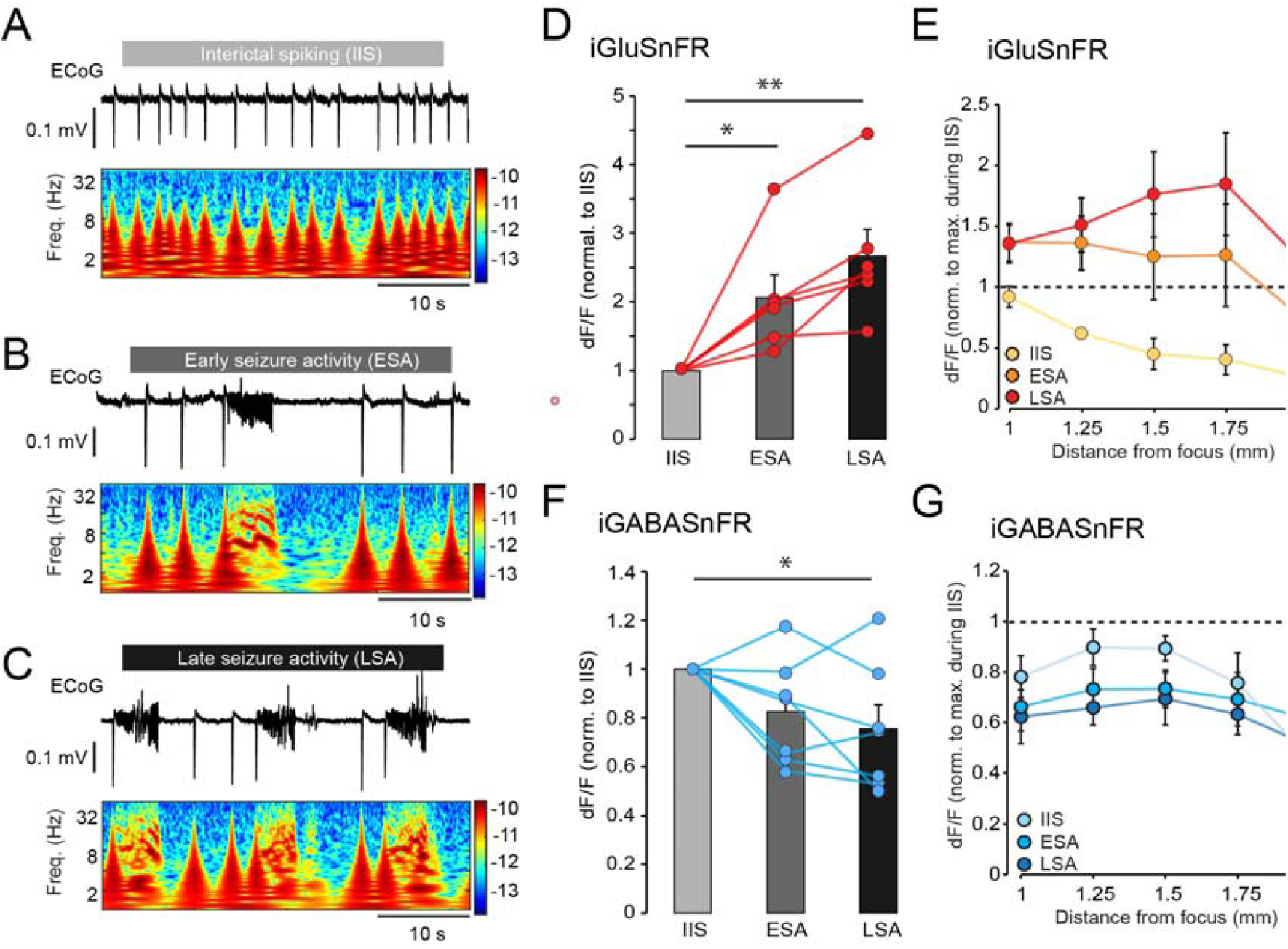
Increase of glutamate release and decrease of GABA release during evolution from spiking to seizures. (**A**) ECoG wavelet spectrogram during interictal spiking activity (IIS), and early (**B**) and late seizure activity (**C**) (ESA and LSA). (**D**) Peak amplitude of iGluSnFR transients during IIS, ESA and LSA. Note the rapid increase of glutamate release as seizures start to appear (n=6 mice). (**E**) iGluSnFR transients increased in amplitude and broadened out as seizures took over from spikes (n=6 mice). (**F**) Same as D but for iGABASnFR. Note the gradual decrease of GABA release (n=7 mice). (**G**) iGABASnFR transients decreased in amplitude and flattened out (n=5 mice during IIS and ESA, and 4 mice during LSA). Error bars correspond to s.e.m..** p<0.01, *p<0.05.

Taken together, these results reveal a progressive erosion of inhibition close to the focus, as IIDs gradually give way to intermittent seizures, and show that this is mediated at least in part by a decrease in GABA release in an annulus surrounding the focus. Conversely, the evolution of epileptiform activity is associated with an increase in the spatial extent of spike-associated glutamate release, as expected from the progressive recruitment of an increasing population of excitatory neurons, permitted by a weakening of inhibitory restraint.

### Interneuron fatigue is sufficient to account for the transition from interictal spikes to seizures

In order to better understand whether a failure of GABA release could account for the main findings of the present study, we built a computational neural field model, representing a two-dimensional sheet of excitatory and inhibitory neurons, interacting according to the connectivity rules indicated in **Fig. 7A**. A small central area of exogenous excitation was incorporated to mimic the microinjection of a chemoconvulsant (**Fig. 7A**). This excitatory drive led to rhythmic bursts of activity, with near-synchronous glutamate and GABA transients (**Fig. 7B**) similar to those observed to accompany experimental interictal spiking (**Fig. 2A, B**). The simulated inhibitory neuronal population was designed to undergo gradual ‘fatigue’, representing failure of either recruitment or GABA release, after prolonged repetitive activity (**Fig. 7C, D**). Consistent with experimental findings (**Fig. 6E and G**), the GABA transients close to the focus diminished in peak amplitude, while the glutamate transients increased in spatial extent (**Fig. 7C-D**). At around the 41s mark, the activity escaped from local inhibitory restraint and propagated in a self-sustained manner. This simulated seizure invaded surrounding territories, where the inhibitory population was not fatigued, in spiral waves (**Fig. 7D** and sup**plementary video 1**).

**Figure 7:**
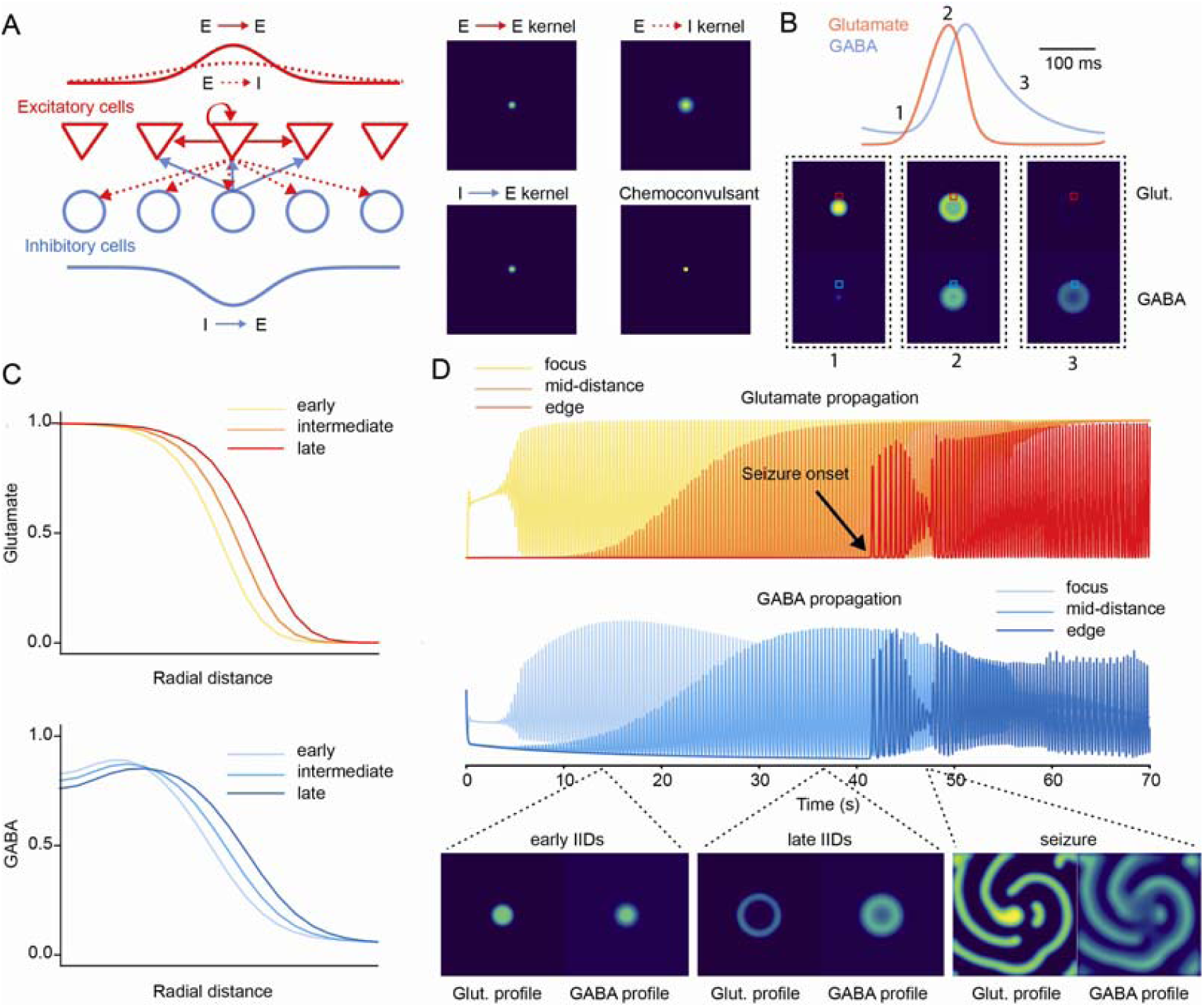
Mean field computational model of seizure initiation: **(A)** Schematic representation of the connectivity between excitatory (E) and inhibitory (I) populations. E to I and I to E connections have similar distributions, although the E to I connectivity is set up with a broader spatial extent as determined by a larger σ parameter on the two-dimensional gaussian connectivity kernel. The chemoconvulsant action was simulated as a small area of external excitatory input to both populations in the centre of the simulated patch of tissue. **(B)** Example glutamate and GABA spatio-temporal profiles during interictal activity (red and blue squares indicate the locations where the glutamate and GABA signals where extracted). **(C)** Peak amplitude of interictal glutamate and GABA waveforms plotted against distance from the simulated chemoconvulsant injection site. Three profiles are shown for spikes progressively closer to the transition to seizure (early, intermediate, and late). **(D)** Temporal evolution of glutamate and GABA release over time at the focus, mid-distance and edge of the simulated patch of tissue. The interictal spiking activity slowly invades neighbouring areas as the central inhibitory populations become fatigued. At around the 41s mark the interictal activity escapes and propagates in a self-sustained manner in new areas as yet unaffected by inhibitory population fatigue (black arrow: seizure onset). (Lower panel: spatial still frames of glutamate (left) and GABA (right) release at three timepoints, as interictal activity gives way to a seizure.

## Discussion

The present study reveals strikingly different profiles of extracellular GABA and glutamate during interictal discharges. Whilst glutamate transients are maximal near the focus and propagate centrifugally, GABA transients are maximal approximately 1.5 mm away. This is consistent with an inhibitory annulus surrounding the focus, as expected from recruitment of feed-forward interneurons confining excessive activity and preventing runaway excitation. IIDs detected electrophysiologically are superficially indistinguishable from sentinel spikes accompanying seizures. However, GABA- and glutamate-dependent fluorescence transients at the onset of seizures are smaller and bigger respectively, providing an explanation for the failure of feed-forward inhibition to constrain epileptiform activity. As interictal spiking gradually gives way to seizures, GABA transients further diminish, consistent with a progressive erosion of inhibitory restraint.

Prior to this study, the experimental evidence for failure of inhibitory restraint has come from electrophysiological recordings and Ca^2+^ imaging, both of which have limitations. Electrophysiology samples the cortex sparsely, although recently developed high-density probes are beginning to shed light on the activity of multiple neurons in a small area ^32^. Identifying action potentials generated by individual neurons through the evolution of a seizure is however complicated by poor signal to noise ratios and distortion of the extracellular fields and spike waveforms, and the full potential of such high-density probes remains to be determined. Ca^2+^ imaging, on the other hand, generally has a slow temporal resolution dictated by the off-rate of fluorescent indicators ^33^, and has not yet been multiplexed during seizures to allow simultaneous imaging of different neuronal subtypes. Recently optimized voltage indicators show promise because of their fast kinetics and improved dynamic range ^34, 35^, but have yet to be applied to understand seizure mechanisms. Importantly, intracellular Ca^2+^ elevation and neuronal depolarization can occur in the absence of neurotransmitter release if neurons enter a state of depolarization block. Fluorescence imaging using iGluSnFR and iGABASnFR fills an important gap in that it provides a direct measure of the interplay of excitatory and inhibitory signalling.

Previous single cell electrophysiology studies have shown that pyramidal neurons receive predominantly synchronous GABAergic currents during IIDs elicited by chemoconvulsants and/or pro-convulsant perfusion solutions. This has been reported both in brain slices obtained from rodents ^6, 10, 18, 20^ and in tissue resected from epileptic patients ^4, 12^. Synchronous Ca^2+^ transients in interneurons also occur during interictal spikes in a chronic epilepsy model^5^. Our imaging data extend these findings by revealing intense GABA release during IIDs, with iGABASnFR fluorescence time courses suggestive of slow neurotransmitter clearance, but also show quasi-synchronous glutamate release. A striking difference, however, is that the glutamate transients diminished in peak amplitude with increasing distance from the focus, whilst GABA transients were maximal ∼1.5 mm from the site of injection. In addition, the iGluSnFR and iGABASnFR fluorescence waves accompanying IIDs propagated at different velocities and in opposite directions: rapidly and centrifugally in the case of glutamate and more slowly and centripetally in the case of GABA. A caveat should be stressed, however: our experimental design did not allow us to image the neurotransmitters at the core of the ictal focus. Nevertheless, the iGluSnFR and iGABASnFR fluorescence recordings provide direct evidence for surround inhibition ^36^ constraining epileptiform activity inside the focus. It also confirms that a similar inhibitory restraint phenomenon takes place during IIDs at the focus as observed during ictal propagation in animal models and epileptic patients ^7, 9^. Notably, the glutamate waves observed in our study propagated at a similar speed (∼100 mm/s) as the LFP in some patients with epilepsy in whom grids of electrodes were used to monitor cortical seizures ^9^. The GABA wave was 2 to 3 times slower. A simple model to account for the results is that a ‘detonation’ of glutamatergic neurons at the focus recruits feedforward interneurons, resulting in an annulus of GABA release. Depolarization block of proximal interneurons ^20^ could then explain why less GABA is released closer to the focus.

A striking finding in the present study was that sentinel spikes heralding seizure onset were accompanied by smaller iGABASnFR transients and bigger glutamate transients than IIDs. A previous study using Ca^2+^ fluorescence imaging in pyramidal neurons in a similar awake head-fixed model failed to distinguish between IIDs and seizures during the first 300 ms ^11^. A partial failure of GABA release is a strong candidate explanation for why inhibitory restraint collapses intermittently. Another *in vitro* study using low [Mg^2+^] to evoke epileptiform activity reported a gradual decrease of synchronous GABA_A_ receptor-mediated currents, and an increase in glutamatergic currents, in pyramidal cells and interneurons starting between 3 and 5 IIDs prior to seizure onset ^18^. This provides a remarkable match with our observations in the five animals where we were able to capture a sufficient number of events (**Fig. 5**). These data strongly suggest that a decrease in GABA release and an increase in glutamate are causally upstream of the transition of IIDs to seizures. We also observed a gradual decrease of GABA release and a broadening of the glutamate waves with each event (whether IID or sentinel spike) accompanying the evolution of seizure activity over minutes. This pattern is in accord with a progressive loss of inhibitory barrages seen in acute slices while the frequency of ictal events increased ^10^.

A simple mean field model reproduced the main experimental findings, and only required the inhibitory population to undergo fatigue upon repetitive firing for rhythmic spiking to give way to a seizure. Why does GABA release partially collapse with the discharges leading up to seizures? Although depolarization block of interneurons ^19, 20^ could account for this observation, it could, in principle, also be explained by vesicle depletion ^18^ or presynaptic inhibition of GABAergic terminals ^17, 37^. These phenomena are not mutually exclusive, and all three could, in principle, occur, especially when taking into account the diversity of interneurons. However, vesicle depletion and presynaptic autoinhibition mechanisms typically require repeated spiking activity above 10 Hz ^17, 38, 39^, and such short-term depression would be expected to have faded before the next interictal spike. Supported by a recent *in vitro* study^40^, we suggest that intense depolarization, preventing a fraction of interneurons from firing, is the most plausible explanation for the partial collapse of GABA release at seizure onset. Although this mechanism has the attraction that it also explains why GABA release is maximal in an annulus around the ictal core, it remains to be directly tested *in vivo*.

A limitation of our experimental design is that, because we imaged the superficial cortical layers, the conclusions relate to glutamate and GABA release onto apical dendrites of pyramidal neurons. Future investigations to study spatiotemporal profiles of glutamate and GABA during epileptic activity across cortical layers would be of great interest ^23^. In addition, identifying the sources of the GABA transients would shed light on whether the partial collapse of GABA release is specific to a particular interneuron subtype. Layer 1 includes at least 4 different types of interneurons ^41^ as well as axonal terminals from somatostatin-positive interneurons ^42^, and so there are several candidates. A further caveat is that the present study used acute chemoconvulsant models. However, the ability to model focal cortical dysplasia, which is associated with spontaneous seizures ^43, 44^, as well as open-source miniaturized microscopes ^45, 46^, may allow glutamate and GABA release to be studied in chronic epilepsy models.

To conclude, we have provided direct evidence that the gradual collapse of inhibitory restraint in awake animals is associated with a progressive decrease in GABA release. Future investigation of the precise mechanisms underlying this failure, and the interneurons responsible, may help to design new strategies to prevent epileptiform activity from invading new territories.

## Methods

### Animals

All experimental procedures on animals were conducted according to the UK Animals Scientific Procedures Act (1986). Experiments were performed at University College London, under personal and project licenses released by the Home Office following appropriate ethics review. *CaMKII-Cre* (B6.Cg-Tg(*CaMK2a*-Cre)T29-1Stl/J, stock no.: 005359, Jackson laboratory) or wild-type C57BL/6 (Harlan) mice were used, with fluorescent reporters expressed using adeno-associated viral vectors carrying either iGluSnFR or iGABASnFR constructs. Animals were housed on 12□h/12□h dark/light cycle, and food and water were given ad libitum.

### Surgery and viral injection

Animals of both sexes (P30-50) were deeply anaesthetized in an induction chamber with 4-5% isoflurane. They were then transferred into a stereotaxic frame (David Kopf Instruments Ltd., USA) and maintained under anaesthesia with 1.5-2.5% isoflurane. Buprenorphine (0.02 mg/kg) and Metacam (0.1 mg/kg) were injected subcutaneously before starting any surgery, and dexamethasone (2 mg/kg) was administered by intramuscular injection before performing the craniotomy. In order to image iGABASnFR or iGluSnFR, animals were subjected to two procedures.

First, mice were injected in layer 5 of the visual cortex using a microinjection pump (WPI Ltd., USA), 5□µl Hamilton syringe (Esslab Ltd., UK), and a 33 gauge needle (Esslab Ltd., UK) (injection volume per site: 350□nL, rate: 100□nL/min) with either AAV2/5.*hSyn*.Flex.iGluSnFR.WPRE.SV40 (titre: 2.5×10^13^ GC per ml, Penn Vector Core, Fig. 2-6 and Fig. S2 and S5) or

AAV2/1.*hSyn*.Flex..iGABASnFR.F102G.mRuby3 (titre: 1.0×10^13^ GC per ml, Janelia Farm, Fig. 2-6 and Fig. S1, S2 and S5).

In some cases, animals were injected with

AAV2/1.*hSyn*.Flex.iGABASnFR.F102G (titre: 1.0×10^13^ GC per ml, Janelia Farm, Fig. 2-6 and Fig. S1, S2 and S5), a new GABASnFR2 variant AAV5/1.*hSyn*.iGABASnFR2^47^ (titre:1.3×10^13^, Janelia Farm, Fig. 5)

or a mixture (1:1) of AAV2/1.*hSyn*-SF-Venus-iGluSnFR.A184V (titre: 9.5×10^12^ GC per ml Janelia Farm) and AAV2/1.*hSyn*.Flex.iGABASnFR.F102G.mRuby3 (titre: 1.0×10^13^ GC per ml, Janelia Farm, Fig. S1). After 10 min, the needle was retracted slowly to the surface, and the animal was sutured and given 0.5-1 mL sterile saline subcutaneously. Mice were checked daily for several days post-surgery.

Three to four weeks post-injection, animals were chronically implanted with a head-post (Model 9, Neurotar Ltd), and a cranial window of 2 mm was performed using a punch hole (Kain medical) in the vicinity of the viral injection site. The craniotomy was then covered with a coverslip (3 mm) which was glued in place. Posterior to the cranial window a small burr hole was drilled for subsequent injection of chemoconvulsant drugs, and a stainless-steel wire (0.075 mm, Advent Ltd) was implanted at the cortical surface nearby to record the ECoG (**Fig. 1**). A reference electrode was implanted in the contralateral frontal lobe. The head-post, electrodes and connectors were secured by opaque dental cement, and silicone elastomer sealant (Kwik-Cast, World Precision Instruments) was used to cover and seal the entire chamber area in order to protect the imaging window and site of injection (**Fig. 1**). Animals were then allowed to recover before habituation in the head-fixed system (Neurotar Ltd).

### Simultaneous imaging and ECoG recordings

Imaging data were acquired on an FV1200MPE multi-photon laser scanning microscope with a X25 immersion objective (XLPLN25XWMP2, Olympus), and adapted to allow a head-fixed system (Neurotar Ltd). iGABASnFR and iGluSnFR were excited at 920 nm with a Ti-sapphire Chameleon Ultra pulsed laser (Coherent), and emission fluorescence was collected via 3 photomultipliers and filters (PMT 1: 450-500 nm; PMT 2: 515-560 nm; PMT 3: 590-650 nm). iGluSnFR and iGABASnFR imaging was performed by using a spiral line scan at 40-60Hz at 320 × 320 pixel (512 × 512 *μ*m, **Fig. 1**) and laser power was kept constant within each experiment. Each imaging acquisition lasted ∼20 sec during the initial phase and was increased to ∼40 sec after detection of the first seizure. Three overlapping imaging loci between 1 mm and 2 mm from the injection site, covering approximatively 1 mm in the antero-posterior axis, were selected before starting each recording session (**Fig. 1**, **Fig. 3-S1**). The centre points of each imaging locus, and distances from the chemoconvulsant injection site, were recorded from the X, Y coordinates given by a motorized platform (Scientifica) holding the head clamp (Neurotar Ltd). Different areas were repeatedly imaged in a pseudorandom order (**Fig. 3_S1**).

The ECoG signal was acquired during the entire experiment via a MultiClamp 700B amplifier (Molecular Devices) and time-stamped with the imaging data by continuously recording the TTL output signal from the scanning-head. ECoG data were bandpass-filtered from 0.1 to 300□Hz, digitized at 1□kHz and recorded using WinEDR open source software (Strathclyde University, UK).

### Acute experiment protocol

After three or four days of habituation to the head-restraint system, animals were lightly anesthetized (1.5% isoflurane) and placed in a stereotaxic frame (Kopf Instruments) for the chemoconvulsant injection. After removing the silicone membrane covering the chamber and inspection of the craniotomy, a Hamilton syringe needle (33 gauge) was inserted into the brain via the injection site hole to a depth of 500-600 µm below the pia (corresponding to layer 5, **Fig. 1**). Either picrotoxin (PTX, 10 mM, 50-100nL) or pilocarpine (3.5 M, 150-300nL) diluted in sterile saline was injected at 100 nL/min, and the needle was removed ∼100 seconds later. The animal was then transferred to the imaging set-up to start the recording session. During the session, animals were checked intermittently for signs of distress and given water via a 0.5 mL syringe. Typically, a recording session lasted up to 30 – 40 min following the chemoconvulsant injection. If the animal was still seizing at the end of the session, it was given diazepam (10 mg/kg) before returning it to its home-cage. All animals were monitored for several hours after each imaging session. Most data were acquired during a single imaging session. However, in a animals where the chemoconvulsants failed to induce seizures, a second session was performed a few days later.

### Image analysis

Imaging data were initially screened using ImageJ software, and the fluorescence timecourses from each pixel extracted for further analysis in MATLAB (The MathWorks Inc.). All analyses were performed using custom-code MATLAB or Python scripts. Briefly, the average fluorescence signal from each scan line was aligned to the ECoG traces by using the recorded TTL output pulse from the scanning system and each fluorescence point was placed at the centre of the TTL pulse in the electrophysiological recordings. For each trace, *Δ*F/F was calculated by taking the baseline F as the median fluorescence during the entire imaging acquisition (either 20s or 40s, see above).

Average fluorescence signals were aligned to the onset of interictal spikes or sentinel spikes (for seizures) as determined from the electrophysiological trace. To estimate the decay time constant of the iGABASnFR and iGluSnFR signals during interictal spikes, a single monoexponential curve was fitted (**Fig. 2**) from the peak to 1s after the peak. Measurement of the area under the curve (AUC) of the iGABASnFR signal associated with interictal spikes or sentinel spikes was restricted to the first 300 ms to avoid later deflections of the ECoG (**Fig. 6**). In this case, all fluorescent transients were collected at ∼1.5 - 2.5 mm from the focus.

To unmix the fluorescence signals arising from simultaneous imaging of iGABASnFR and iGluSnFR signals (**Fig. 2-S1**), we first established, in separate experiments, the fluorescence profile of each probe alone collected via the three PMTs (**Fig. 2_S1 A**). This allowed us to determine the proportion of fluorescence signal (bleedthrough coefficients A and B) detected by each PMT during interictal spikes for iGABASnFR and a Venus (yellow) version of iGluSnFR with our filter sets. We found that the relative Venus-iGluSnFR signals acquired via PMT_1_, PMT_2_ and PMT_3_ were ∼0:1:0.25, and the relative iGABASnFR signals were ∼1:1:0. We then used these profiles to estimate the contribution of both probes expressed simultaneously, and noise, to the fluorescence signals collected via each PMT as follows:

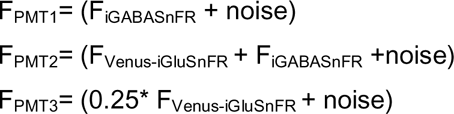

From these formulae, we then separated the Venus-iGluSnFR signal from iGABASnFR fluorescence trace as follows:

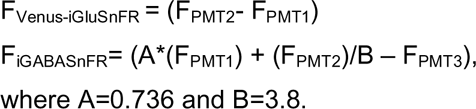

To determine the direction of propagation and velocity of fluorescence signals, we first re-aligned the fluorescence timecourse for each pixel to compensate for the inter-pixel interval (10 µs). We then estimated the time at which regions opposite one another in the imaged area (equidistant from its centre, ∼1.5 mm from the focus) reached 50% of maximal fluorescence intensity during each IIS (T_50_). A vector in the X-Y plane passing through the opposite regions was rotated through 360*°*, and the vector angle that yielded the steepest correlation between T_50_ and the distance along the vector defined the direction of propagation of the neurotransmitter wave (**Fig. 4, S1**). This procedure also allowed us to estimate the speed of propagation of the interictal fluorescence transients.

### Computational model

The neural field model used here is adapted from Liou et al 2020^48^. A patch of tissue was modelled with an excitatory and an inhibitory population at every point in space. The dynamics of the average membrane voltage, *V*, of each population is dictated by:

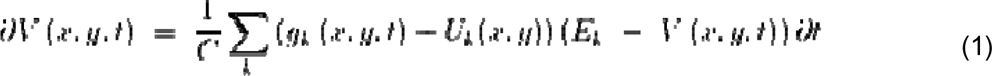

Where *C*, is the average cell capacitance and *g_k_* and *E_k_* are, respectively, the conductance and reversal potential of the channel family *k*. *Uk* is an exogenous input to the channel family *k*. There is a time independent leak conductance and two time-dependent conductances: glutamatergic and GABAergic. The dynamics of the latter two conductances are dictated by:

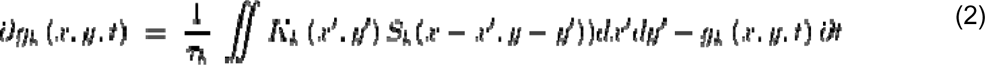

Where *K_k_* is the connectivity kernel to the target population from the glutamatergic or GABAergic populations, whose firing rates are represented by *S_k_.* The firing rates of each population are computed via an instantaneous sigmoid function applied to their average membrane voltage:

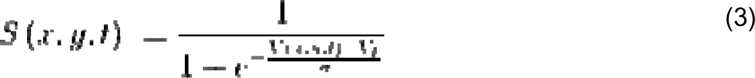

Where *V_t_* represents the firing threshold voltage for the given population, and σ is usually interpreted as the dispersion of membrane voltages across the population. In this work, the modelling of the inhibitory population fatigue mechanism diverged from Liou et al (2020) and used a simplified adaptation of the firing threshold *V_t_* according to a first order relaxation mechanism, following the equation:

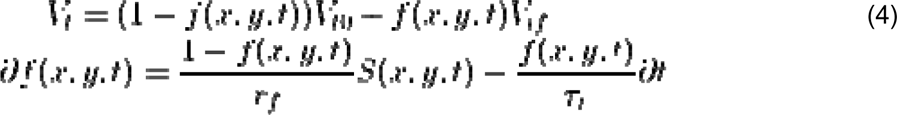

Where *f* is the proportion of fatigued neurons, *V_t0_* is the normal firing threshold of the population and *V_tf_*is the firing threshold the population approaches when all neurons are fatigued. The parameter *r_f_* is the inverse ratio of neurons that become fatigued when firing, and τr is their recovery time constant.

The schematic connectivity of the modelled patch of tissue is presented in **Fig. 7A** and the parameters for each of the population are shown in the supplementary information **Supplementary Table 1**.

The python code for the simulations is available at https://github.com/KullmannLab/Shimoda2023.

### Chemicals

Picrotoxin and pilocarpine hydrochloride were purchased from Tocris (CAS number: 124-87-8) and Sigma-Aldrich (CAS number: 54-71-7) respectively and diluted on the day of the experiment in sterile saline.

### Statistics

All statistical analyses were performed either using OriginPro software or in MATLAB. Deviation from normality was tested for each data set and parametric or non-parametric tests were used appropriately. Paired and unpaired t-tests were always two-tailed. To estimate the direction of propagation of the iGABASnFR and iGluSnFR signals, angles were averaged for each animal and compared using a Parametric Watson-Williams multi-sample test. *N* values used in statistical analysis represent the number of animals in each experimental group. Data are shown as mean ± s.e.m.. Statistical differences were considered significant at p < 0.05.

## Acknowledgements

We are grateful to members of the Experimental Epilepsy Group at the UCL Queen Square Institute of Neurology for advice and technical assistance. This work was supported by Epilepsy Research UK, the Medical Research Council and The Wellcome Trust. V.M. is the recipient of an Emerging Leader Fellowship from Epilepsy Research UK (F1901). Y.S. received funding from the Japan Epilepsy Research Foundation.

## Supplementary Figures

**Figure S1:**
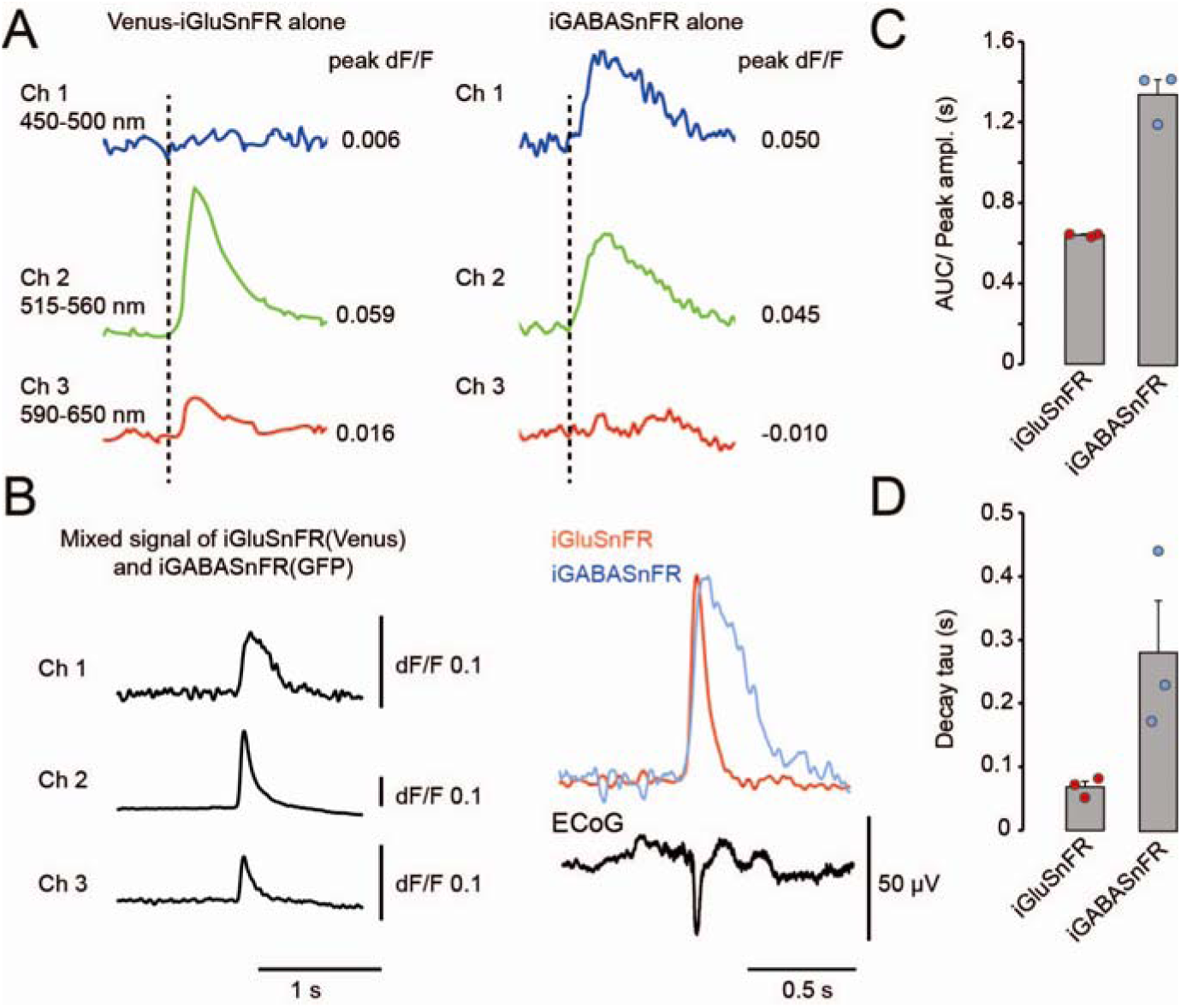
Simultaneous imaging of iGABASnFR and iGluSnFR during interictal discharges. (**A**) Separate imaging of Venus-iGluSnFR (left panel) or iGABASnFR (right panel) fluorescence signal during interictal spikes via the three PMT channels, used to estimate coefficients necessary to unmix signals from both fluorophores during dual imaging. Averaged fluorescence traces (Venus-iGluSnFR, 28 events; iGABASnFR, 12 events) were aligned to IIS onset (dashed line). (**B**) Averaged fluorescence traces (23 events) via the three PMT channels from both sensors imaged simultaneously (left panel) and result of unmixing (right panel), together with simultaneously acquired ECoG trace. (**C**) Average Area Under the Curve (AUC) divided by the peak amplitude for unmixed iGABASnFR and iGluSnFR signals. (**D**) Same as C but for the decay time constant estimated from peak to 1s post-peak. Consistent with Figure 2, GABA transients were almost 3-fold longer than glutamate transients (n=3 animals, Error bars correspond to s.e.m.).

**Figure S2:**
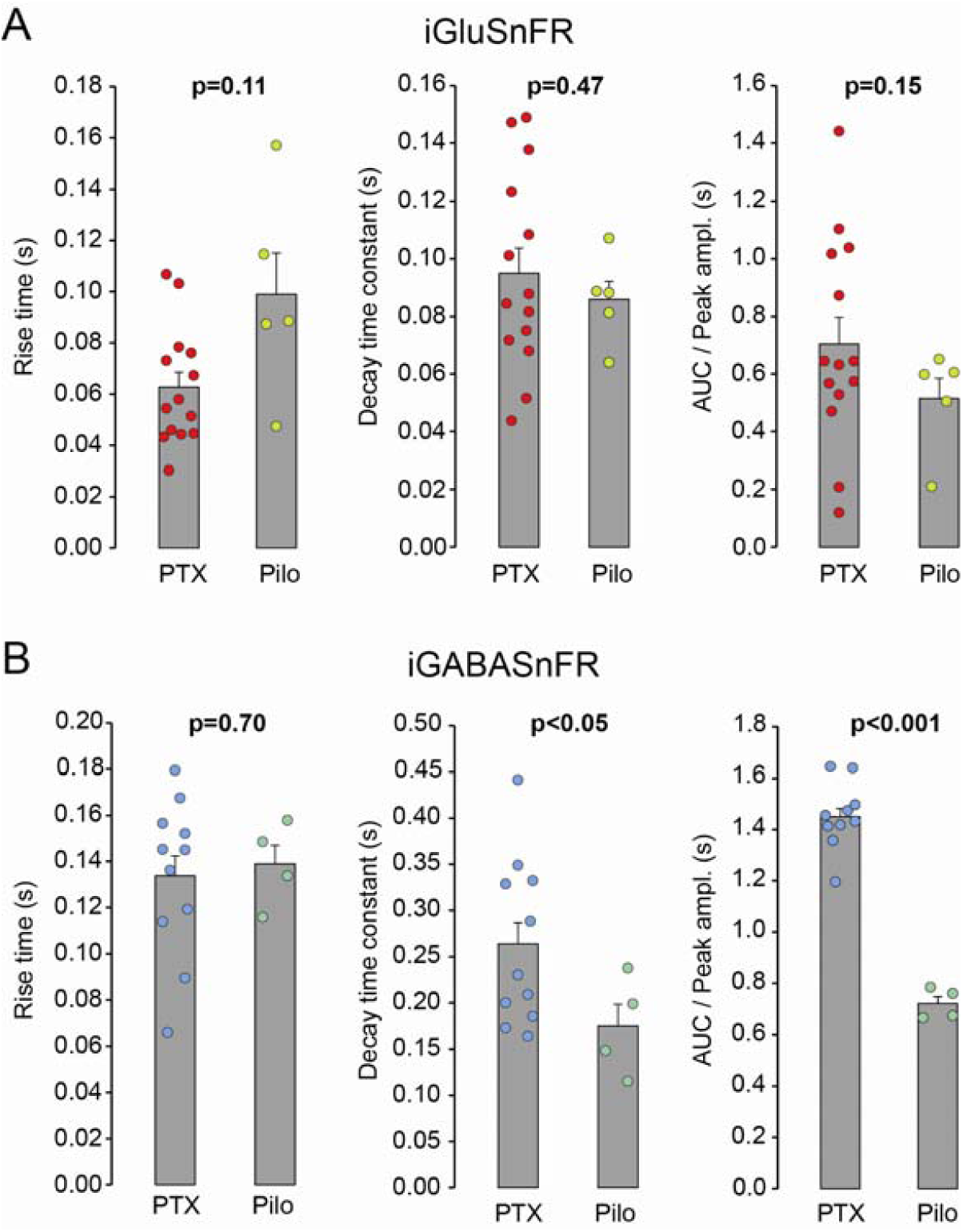
iGABASnFR and iGluSnFR fluorescence transients are broadly similar between the picrotoxin and the pilocarpine models. (**A**) Rise time, decay time constant, and area under the curve (AUC) normalized by the peak amplitude, for iGluSnFR transients associated with IIDs in the PTX (n=14) and Pilo (n=5) models. (B) Similar analysis for iGABASnFR. Note that the decay time constant and AUC were significantly longer in response to PTX (n=11) than Pilo (n=4). Unpaired t-test.

**Figure S3:**
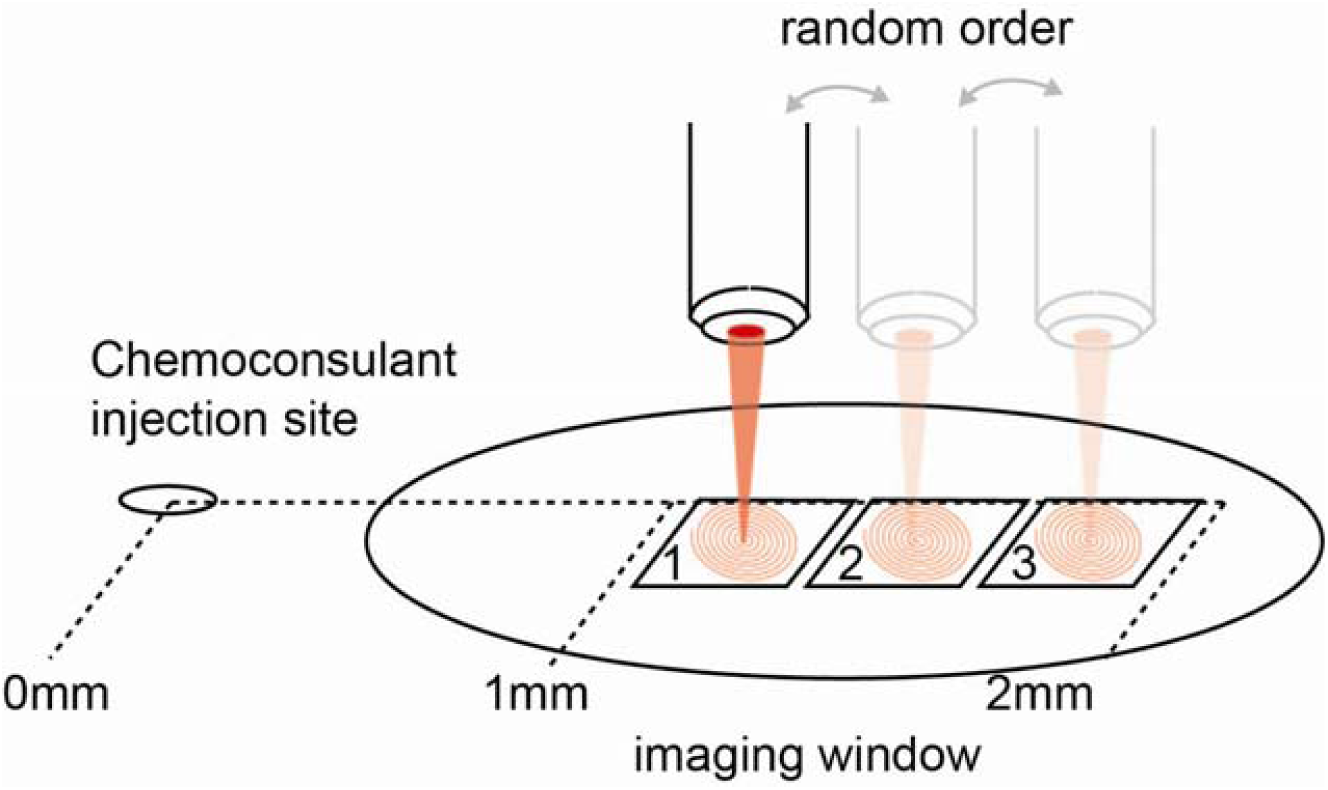
Glutamate and GABA transients vary with distance from the focus. (**A**) Experimental design used to image at different locations during the interictal spiking period. At least 3 regions of interest were pre-selected before chemoconvulsant injection and then imaged in a pseudorandomized order.

**Figure S4:**
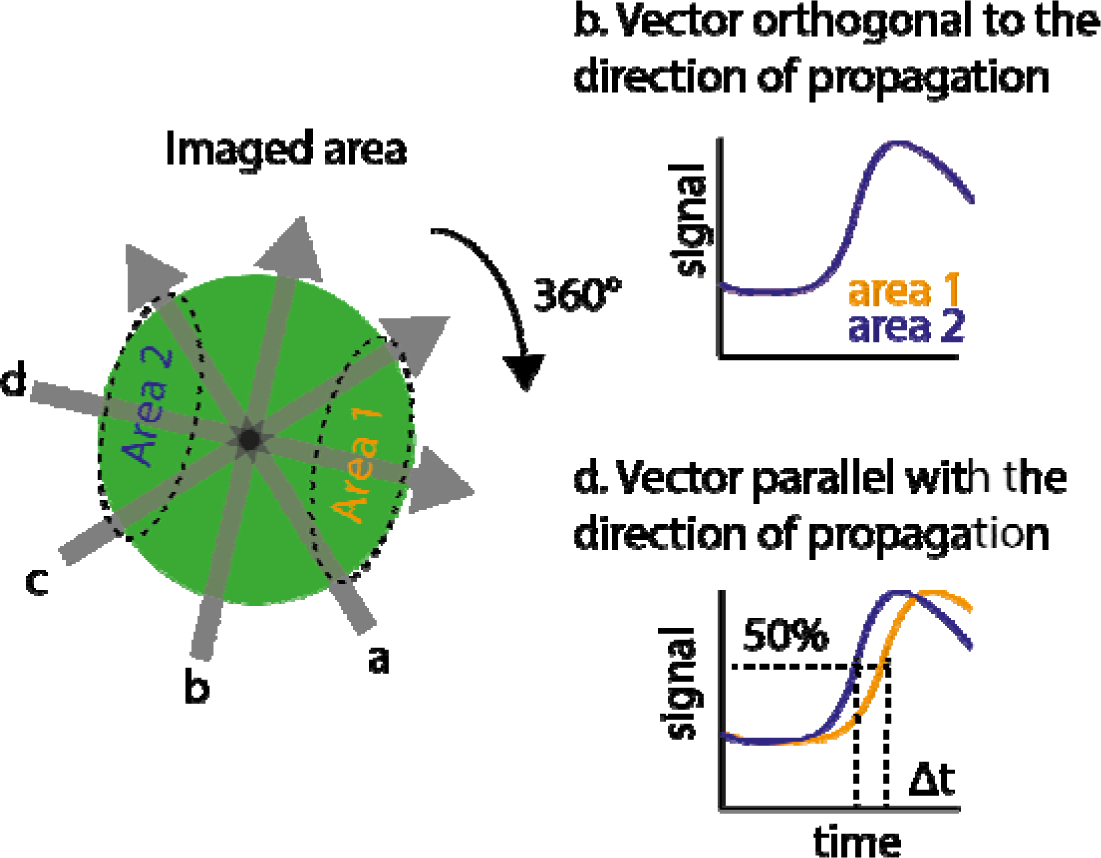
Method to determine the direction and propagation speed of the glutamate and GABA waves during interictal discharges.

**Figure S5:**
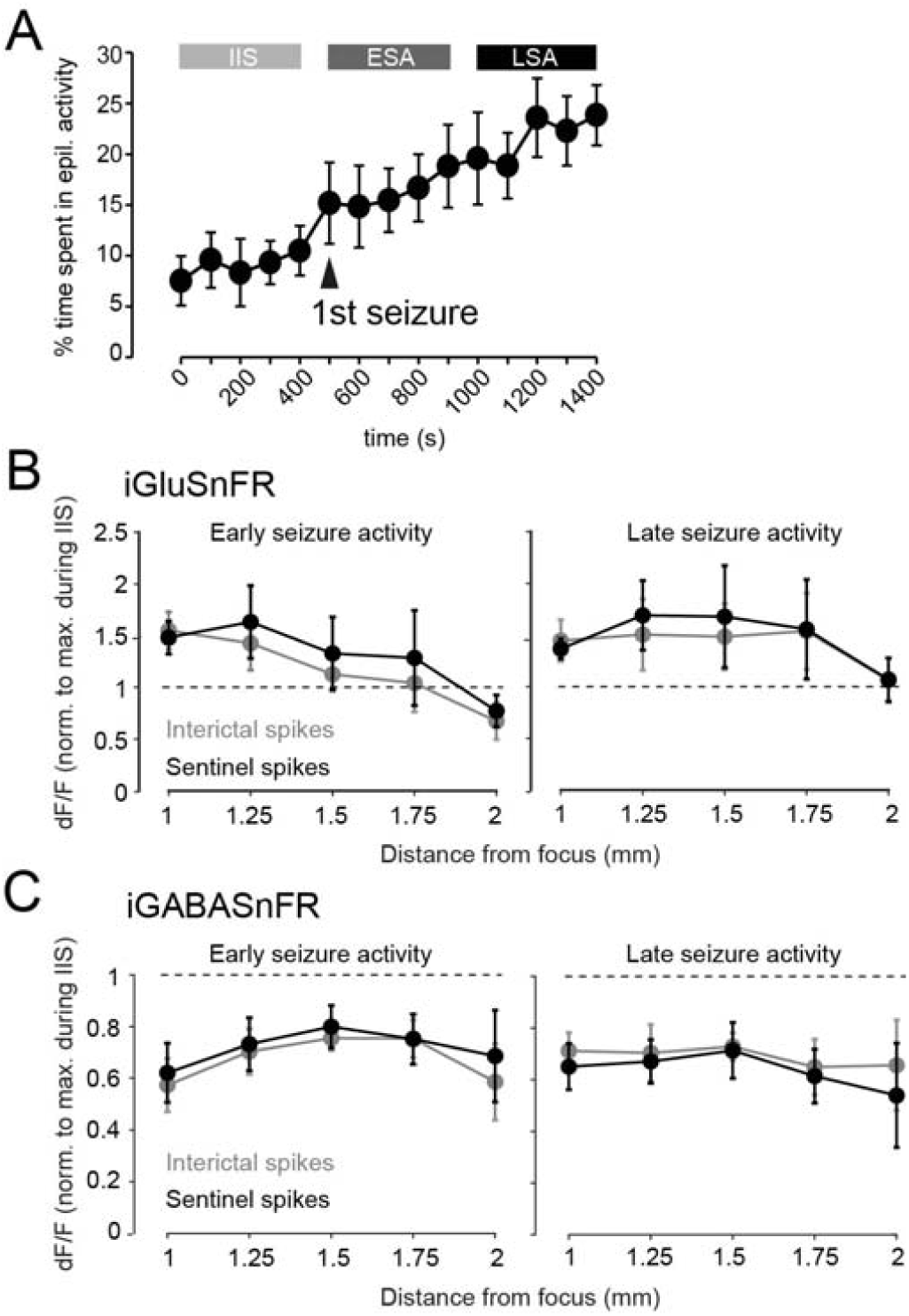
Interictal and sentinel spikes during the early and late seizure phases. (**A**) Percentage of time spent in epileptiform activity (IIDs or seizures) over the three periods (IIS, ESA, LSA; n=8 mice). Note the gradual increase with an abrupt transition after the first seizure. (**B**) Normalized peak amplitude of iGluSnFR and iGABASnFR fluorescence transients during interictal and sentinel spikes at different distances from the focus, during ESA and LSA. Transients associated with IISs and sentinel spikes show similar spatial profiles within each phase. (**C**) AUC/peak amplitude plotted against peak amplitude (both normalized by the average values within each experiment) for iGABASnFR. (**C**) Same as (B) for iGluSnFR. Each point represents an individual GABA or glutamate transient during either interictal or sentinel spikes during the seizure activity period. Error bars in (A) correspond to s.e.m..

**Table 1:**
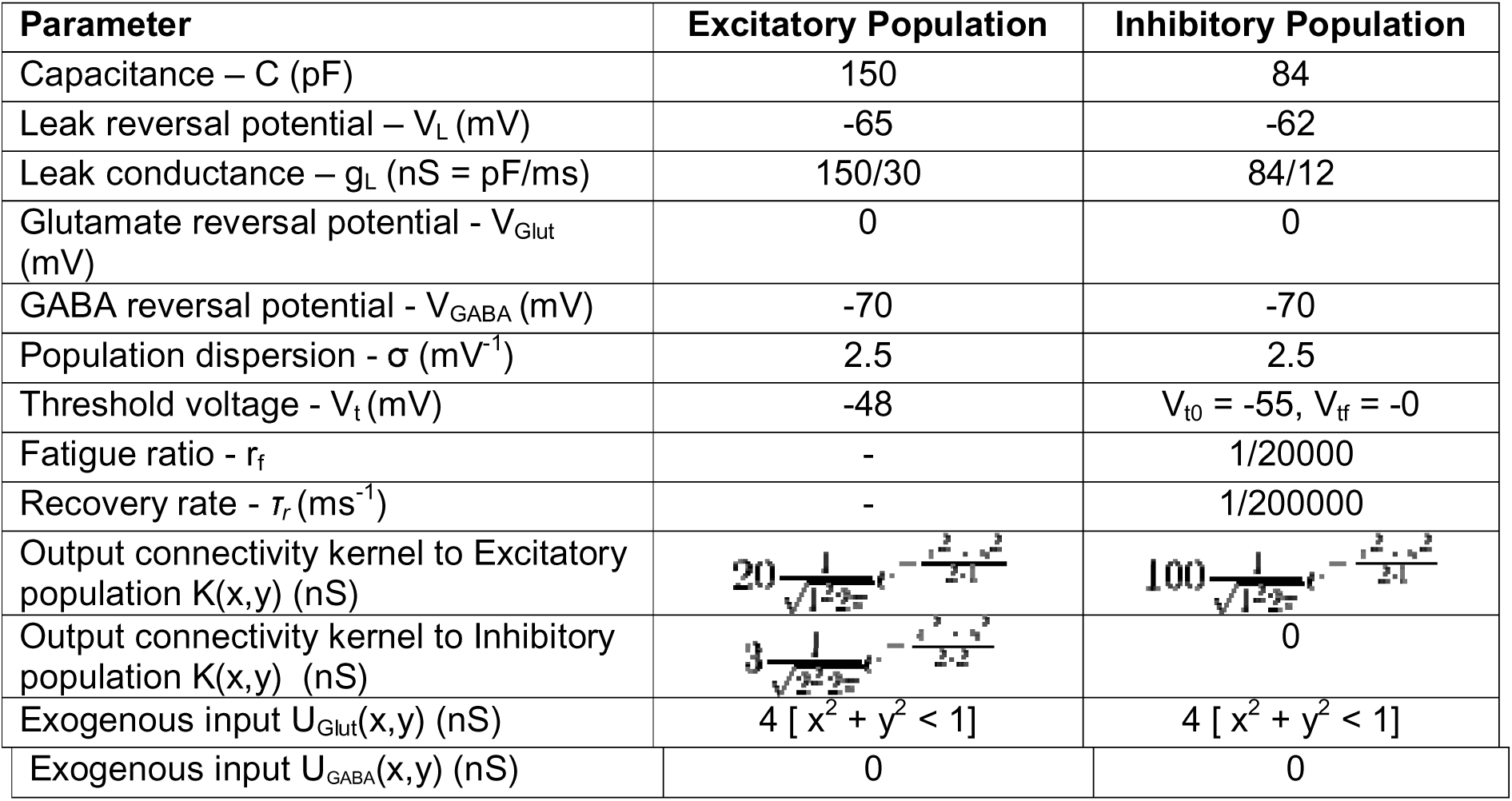
Parameters of the mean field computational model of seizure initiation used in Fig. 7.

## Notes

### Competing Interest Statement

The authors have declared no competing interest.

### Summary of Updates

We have added additional data demonstrating similarity of results obtained with two chemoconvulsant models, and a computational simulation that recapitulates the main experimental findings (Figure 7). We have also included a new Table and supplementary materials including a video of the simulation.

## References

1. de Curtis, M. & Avanzini, G. Interictal spikes in focal epileptogenesis. Progress in Neurobiology 63, 541–567 (2001).

2. Lévesque, M., Salami, P., Shiri, Z. & Avoli, M. Interictal oscillations and focal epileptic disorders. European Journal of Neuroscience 48, 2915–2927 (2018).

3. Bohannon, A. S. & Hablitz, J. J. Optogenetic dissection of roles of specific cortical interneuron subtypes in GABAergic network synchronization. The Journal of Physiology 596, 901–919 (2017).

4. Huberfeld, G. et al. Glutamatergic pre-ictal discharges emerge at the transition to seizure in human epilepsy. Nature Neuroscience 14, 627–634 (2011).

5. Muldoon, S. F. et al. GABAergic inhibition shapes interictal dynamics in awake epileptic mice. Brain 138, 2875–2890 (2015).

6. Trevelyan, A. J., Sussillo, D., Watson, B. O. & Yuste, R. Modular Propagation of Epileptiform Activity: Evidence for an Inhibitory Veto in Neocortex. Journal of Neuroscience 26, 12447–12455 (2006).

7. Liou, J. et al. Role of inhibitory control in modulating focal seizure spread. Brain 141, 2083–2097 (2018).

8. Parrish, R. R., Codadu, N. K., Scott, C. M.-G. & Trevelyan, A. J. Feedforward inhibition ahead of ictal wavefronts is provided by both parvalbumin- and somatostatin-expressing interneurons. The Journal of Physiology 597, 2297–2314 (2019).

9. Schevon, C. A. et al. Evidence of an inhibitory restraint of seizure activity in humans. Nat Commun 3, 1060 (2012).

10. Trevelyan, A. J., Sussillo, D. & Yuste, R. Feedforward Inhibition Contributes to the Control of Epileptiform Propagation Speed. Journal of Neuroscience 27, 3383–3387 (2007).

11. Rossi, L. F., Wykes, R. C., Kullmann, D. M. & Carandini, M. Focal cortical seizures start as standing waves and propagate respecting homotopic connectivity. Nature Communications 8, 217 (2017).

12. Cohen, I. On the Origin of Interictal Activity in Human Temporal Lobe Epilepsy in Vitro. Science 298, 1418–1421 (2002).

13. Pavlov, I., Kaila, K., Kullmann, D. M. & Miles, R. Cortical inhibition, pH and cell excitability in epilepsy: what are optimal targets for antiepileptic interventions? J Physiol 591, 765–774 (2013).

14. Avoli, M., Louvel, J., Kurcewicz, I., Pumain, R. & Barbarosie, M. Extracellular free potassium and calcium during synchronous activity induced by 4-aminopyridine in the juvenile rat hippocampus. The Journal of Physiology 493, 707–717 (1996).

15. Librizzi, L. et al. Interneuronal Network Activity at the Onset of Seizure-Like Events in Entorhinal Cortex Slices. J. Neurosci. 37, 10398–10407 (2017).

16. Viitanen, T., Ruusuvuori, E., Kaila, K. & Voipio, J. The K+–Cl− cotransporter KCC2 promotes GABAergic excitation in the mature rat hippocampus. The Journal of Physiology 588, 1527–1540 (2010).

17. Lei, S. & McBain, C. J. GABAB receptor modulation of excitatory and inhibitory synaptic transmission onto rat CA3 hippocampal interneurons. J Physiol 546, 439–453 (2003).

18. Zhang, Z. J. et al. Transition to Seizure: Ictal Discharge Is Preceded by Exhausted Presynaptic GABA Release in the Hippocampal CA3 Region. Journal of Neuroscience 32, 2499–2512 (2012).

19. Cammarota, M., Losi, G., Chiavegato, A., Zonta, M. & Carmignoto, G. Fast spiking interneuron control of seizure propagation in a cortical slice model of focal epilepsy. J Physiol 591, 807–822 (2013).

20. Karlócai, M. R. et al. Physiological sharp wave-ripples and interictal events in vitro: what’s the difference? Brain 137, 463–485 (2014).

21. Magloire, V., Mercier, M. S., Kullmann, D. M. & Pavlov, I. GABAergic Interneurons in Seizures: Investigating Causality With Optogenetics. Neuroscientist 1073858418805002 (2018) doi:10.1177/1073858418805002.

22. Rossi, L. F., Kullmann, D. M. & Wykes, R. C. The Enlightened Brain: Novel Imaging Methods Focus on Epileptic Networks at Multiple Scales. Front Cell Neurosci 12, 82 (2018).

23. Wenzel, M., Hamm, J. P., Peterka, D. S. & Yuste, R. Reliable and Elastic Propagation of Cortical Seizures In Vivo. Cell Reports 19, 2681–2693 (2017).

24. Wenzel, M., Hamm, J. P., Peterka, D. S. & Yuste, R. Acute focal seizures start as local synchronizations of neuronal ensembles. J. Neurosci. 3176–181 (2019) doi:10.1523/JNEUROSCI.3176-18.2019.

25. Marvin, J. S. et al. A genetically encoded fluorescent sensor for in vivo imaging of GABA. Nat Methods 16, 763–770 (2019).

26. Marvin, J. S. et al. An optimized fluorescent probe for visualizing glutamate neurotransmission. Nature Methods 10, 162–170 (2013).

27. Chiu, C.-S., et al. Number, Density, and Surface/Cytoplasmic Distribution of GABA Transporters at Presynaptic Structures of Knock-In Mice Carrying GABA Transporter Subtype 1–Green Fluorescent Protein Fusions. J. Neurosci. 22, 10251–10266 (2002).

28. Lehre, K. P. & Danbolt, N. C. The Number of Glutamate Transporter Subtype Molecules at Glutamatergic Synapses: Chemical and Stereological Quantification in Young Adult Rat Brain. J Neurosci 18, 8751–8757 (1998).

29. Scimemi, A. Structure, function, and plasticity of GABA transporters. Front Cell Neurosci 8, 161 (2014).

30. Marvin, J. S. et al. Stability, affinity, and chromatic variants of the glutamate sensor iGluSnFR. Nat Methods 15, 936–939 (2018).

31. Magloire, V., Cornford, J., Lieb, A., Kullmann, D. M. & Pavlov, I. KCC2 overexpression prevents the paradoxical seizure-promoting action of somatic inhibition. Nat Commun 10, 1225 (2019).

32. Jun, J. J. et al. Fully integrated silicon probes for high-density recording of neural activity. Nature 551, 232–236 (2017).

33. Kerruth, S., Coates, C., Dürst, C. D., Oertner, T. G. & Török, K. The kinetic mechanisms of fast-decay red-fluorescent genetically encoded calcium indicators. J. Biol. Chem. 294, 3934–3946 (2019).

34. Abdelfattah, A. S. et al. Bright and photostable chemigenetic indicators for extended in vivo voltage imaging. Science 365, 699–704 (2019).

35. Piatkevich, K. D. et al. Population imaging of neural activity in awake behaving mice. Nature 574, 413–417 (2019).

36. Prince, D. A. & Wilder, B. J. Control Mechanisms in Cortical Epileptogenic Foci*: Surround Inhibition. Arch Neurol 16, 194–202 (1967).

37. Overstreet-Wadiche, L. & McBain, C. J. Neurogliaform cells in cortical circuits. Nature Reviews Neuroscience 16, 458–468 (2015).

38. Fernández-Alfonso, T. & Ryan, T. A. The Kinetics of Synaptic Vesicle Pool Depletion at CNS Synaptic Terminals. Neuron 41, 943–953 (2004).

39. Yamashita, M., Kawaguchi, S., Hori, T. & Takahashi, T. Vesicular GABA Uptake Can Be Rate Limiting for Recovery of IPSCs from Synaptic Depression. Cell Reports 22, 3134– 3141 (2018).

40. Călin, A., Ilie, A. S. & Akerman, C. J. Disrupting Epileptiform Activity by Preventing Parvalbumin Interneuron Depolarization Block. J. Neurosci. 41, 9452–9465 (2021).

41. Schuman, B. et al. Four Unique Interneuron Populations Reside in Neocortical Layer 1. J. Neurosci. 39, 125–139 (2019).

42. Tremblay, R., Lee, S. & Rudy, B. GABAergic Interneurons in the Neocortex: From Cellular Properties to Circuits. Neuron 91, 260–292 (2016).

43. Hsieh, L. S. et al. Convulsive seizures from experimental focal cortical dysplasia occur independently of cell misplacement. Nature Communications 7, 1–12 (2016).

44. Barbanoj, A. A. et al. Anti-seizure Gene Therapy for Focal Cortical Dysplasia. 2023.01.09.523292 Preprint at https://doi.org/10.1101/2023.01.09.523292 (2023).

45. Cai, D. J. et al. A shared neural ensemble links distinct contextual memories encoded close in time. Nature 534, 115–118 (2016).

46. Shuman, T. et al. Breakdown of spatial coding and interneuron synchronization in epileptic mice. Nature Neuroscience 23, 229–238 (2020).

47. Kolb, I. et al. Optimization of genetically encoded GABA indicator. (2022) doi:10.25378/janelia.19709311.v3.

48. Liou, J. et al. A model for focal seizure onset, propagation, evolution, and progression. eLife 9, e50927 (2020).

